# Patchy distribution of potato cyst nematodes within single arable fields reveals local disease suppressiveness mediated by disparate microbial communities

**DOI:** 10.1101/2025.02.26.639864

**Authors:** Robbert van Himbeeck, Stefan Geisen, Casper van Schaik, Sven van den Elsen, Roeland Berendsen, Andŕe Bertran, Egbert Schepel, Johannes Helder

## Abstract

Disease suppressiveness is a complex phenomenon that is assumed to be the resultant of actions of local microbial antagonists in soil environments. Exploitation of disease suppressiveness as a tool to manage pathogens is hindered by our poor understanding of this phenomenon. Here we investigated soil microbiome-based suppression of potato cyst nematodes (PCN), and, to this end, four apparently homogeneous potato fields with an unexplained non-homogeneous PCN distribution were selected. We hypothesized that this patchy PCN distribution resulted from local variation in disease suppressiveness. Under controlled greenhouse conditions, we confirmed the suppressiveness of these soils vis-à-vis PCN and soils were gamma-irradiated to corroborate the biotic origin of this suppression. Subsequent DNA-based analysis of the microbial community in the potato rhizosphere revealed suppressiveness-related contrasts in community composition between suppressive and conducive patches. Elevated abundances of fungal (*e.g.*, *Metacordyceps chlamydosporia*) and bacterial (*e.g*., *Pseudomonas fluorescens*) nematode antagonists were positively correlated with PCN suppressive patches. Distinct sets of antagonists were found to be associated with PCN suppression despite of the geographical closeness of the locations under investigation. Our findings confirm the biotic origin of local PCN suppressiveness and reveal that it should be regarded as a superficially similar resultant of a biologically diverse phenomenon.

## 1 Introduction

To restore and protect natural and arable soils, numerous agrochemicals that used to be applied for plant pathogen control are phased out in many global regions (*e.g*., European Union; European Union, 2022). Crop farmers consequently have an increasingly narrow set of control measures at their disposal and especially for soil-borne diseases, novel management strategies are needed. The exploitation of local disease-suppressive potential of the soil microbiome is a promising approach that could contribute to plant-pathogen control (e.g., Expósito et al., 2017). The first description of local disease suppression in soil dates back nearly a century (Henry, 1931), and since that time disease suppressive soils have been scrutinized to isolate and characterize microbial antagonists responsible for this phenomenon. Decades of research has resulted in the discovery of a wide range of natural enemies of plant-pathogenic bacteria (Abo-Elyousr and Hassan, 2021), fungi (Penton et al., 2014; Bubici et al., 2019), and nematodes (Li et al., 2015; Topalovíc et al., 2020). Based on these findings, numerous biological control agents have been developed (*e.g.,* Li et al., 2015; Bubici et al., 2019), but in general, non-native microorganisms show difficulties in establishing in local, highly competitive soil microbial communities. Local microbes have ample competitive competences within their native soil ecosystem. Although it has been shown that stimulating the local pathogen-antagonistic community is feasible, for example through the use of cover crops (Cazzaniga, Belliard, et al., 2025; Cazzaniga, Braat, et al., 2023), our limited understanding of the underlying biological processes (Expósito et al., 2017) currently hampers the broader adoption of this approach as a pathogen management strategy. Characterization and ecological understanding of the microbiome of suppressive soils is essential in harnessing its potential in management strategies. The comparison of microbiomes from suppressive and conducive fields can be complicated if those fields are spatially dispersed, have disparate soil types, or are managed differently. Ideally, one would like to study visually homogeneous fields with an unexplained patchy distribution of a given soil-borne pathogen, while cropping and disease history suggests this pathogen should have been present throughout the field. Such a patchy pathogen distribution could result from hard-to-observe local variations in abiotic conditions or from the recent introduction of the pathogen. However, the success of pathogen invasion might also be caused by local heterogeneity in the composition and functioning of soil microbiomes (Wei et al., 2019; Gu et al., 2022). High-resolution data on the within-field distribution of soil-borne pathogens would be a good starting point for investigating the nature of local disease suppressiveness, but such data are scarce. In this respect, soil-borne pathogens with quarantine status might form an exception as statutory regulations prompt farmers to have their fields sampled on a regular basis and with a relatively high sampling intensity.

Potato cyst nematode (PCN) is the common name for two quarantine pathogens, *Globodera pallida* and *G. rostochiensis*, that are present in virtually all major potato growing regions in the world. These pathogens constitute a major threat to global potato production causing an estimated global yield loss of 9% (Jones et al., 2013). Because of its quarantine status, and given that both seed and consumer potatoes are large export commodities, this soil-borne pathogen is carefully monitored both by plant health authorities and producers. As a result, the distribution of PCN is well-known and numerous distribution maps are available. Although most fields used for potato production are labeled as either non-infested or infested with PCN, high-resolution screenings occasionally reveal a patchy within-field PCN distribution. Particularly these fields are suitable objects to study local suppressiveness vis-à-vis this pathogen. PCN suppressive soils and antagonists have been documented in North-Western Europe (Jones, 2024; Cronin et al., 1997; Velvis and Kamp, 1996), demonstrating the capacity of the region’s soil microbiome to suppress this pathogen. However, PCN suppressive soils are considered to be uncommon (Kerry et al., 2009) and the local soil microbiome of PCN suppressive fields is understudied. Comparison of microbiomes associated with conducive and suppressive patches within single fields might allow us to pinpoint how to manipulate and control suppressive soils.

The aim of this study was to identify within-field differences of PCN suppression and to characterize the associated microbial soil communities. We selected four apparently homogeneous arable fields that showed clear within-field differences in PCN density. We verified whether these patchy PCN distributions had a biotic origin and - upon showing the biotic origin - we investigated whether the virtual absence of PCN was associated with the presence of nematode antagonists. High-throughput DNA sequencing was used to characterize fungal and bacterial rhizosphere communities and to identify potential nematode antagonists. With this experimental setup, we addressed the following research questions: 1a) Can the apparently conducive and suppressive nature of patches within the selected potato production-fields be reproduced under greenhouse conditions? 1b) Do observed within-field differences in PCN infection have a biotic origin?; 2) What assemblages of antagonists are associated with PCN suppression; and 3) how do these assemblages vary among suppressive patches between fields?

## 2 Methods and Materials

### 2.1 Field selection and sampling

To study PCN suppressiveness, visually homogeneous fields with a patchy PCN distribution were identified (Fig. 1A). For this, we mined a proprietary database generated by HLB BV (Wijster, The Netherlands), an agricultural consultancy company. This database comprises high-resolution PCN maps of hundreds of arable fields collected over a range of years. For these maps fields were sampled using an automated soil core collector (every 2 m in rows with a width of 6 or 9 m, and up to 150 m in length). Upon microscopic analyses distribution maps of PCN of entire fields were generated (see figure 1A for example). After consulting the farmers to verify field homogeneity, four PCN patchy-fields were selected in Friesland (a northern province in the Netherlands): field E (farmer 1), G (farmer 2), K (farmer 2), S (farmer 3). The two furthest apart fields were separated by 30 km. The infested and non-infested parts of field K were separated by a narrow ditch. PCN distribution of selected fields was determined recently (*<*3 years). Sections of the field that were infested with PCN were labeled ”putatively conducive”, while non-infested are referred to as ”putatively suppressive”. Patchy fields were sampled at the end of March 2023 (before the main growing season). For each field, five random sampling squares of 9 m^2^ were plotted in both the putatively conducive and suppressive parts, resulting in 10 sampling plots per field. For each plot, 14 soil cores were collected using an auger (,04 cm, 0-25 cm deep). These subsamples were pooled, and soil was mixed thoroughly. Per field four samples of each 100 g (two from each of the putatively conducive and suppressive field sections) per field were collected for chemical analyses (see section 2.2). Putatively conducive field sections were always sampled before putatively suppressive sections to prevent potential contamination with antagonistic microbial taxa. Between sections of a field, all materials were cleaned with 70% ethanol. The soil was transferred to the greenhouse at the day of sampling, and air-dried at ambient temperature.

**Figure 1:**
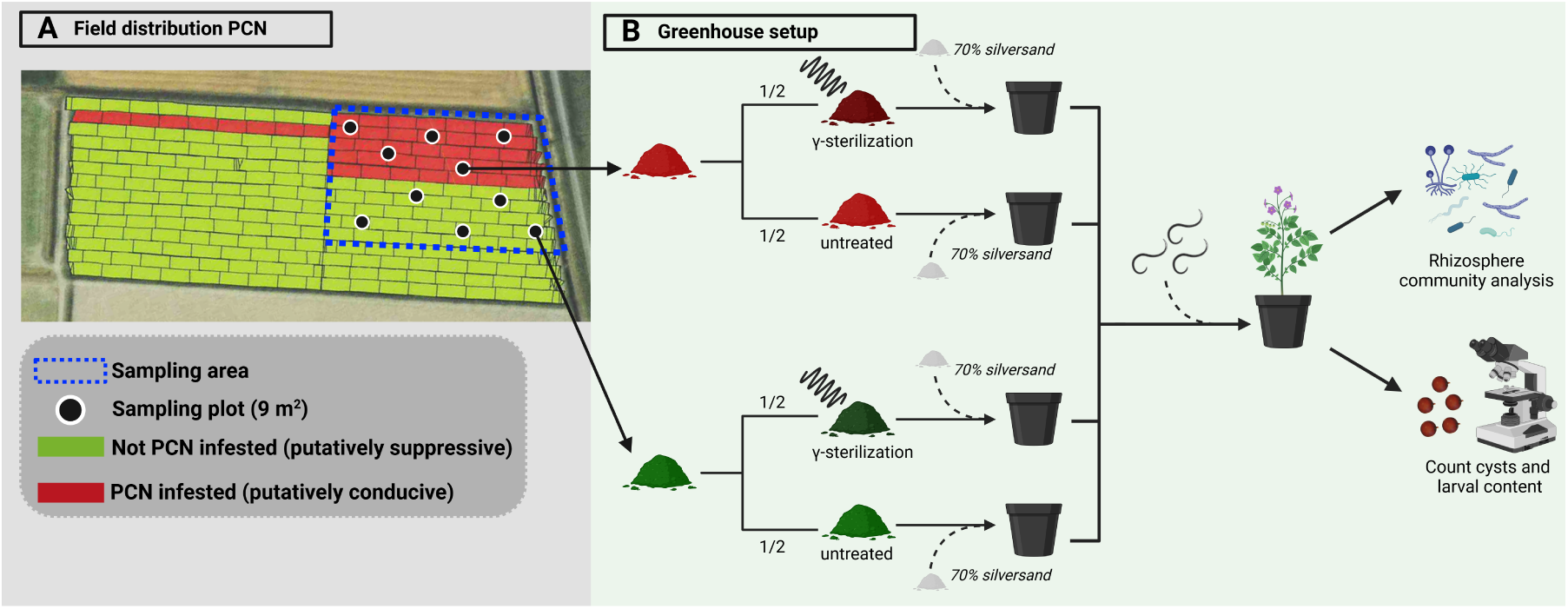
Overview of the experimental setup. A) An example of PCN distribution data and soil sampling scheme. Non-infested and infested field parts identical in size were sampled. In each part, a composite sample was collected from 5 random plots. B) Of each field part, one third of the collected soil was sterilized using *γ*-irradiation, the remaining two thirds were kept untreated. Field soil (30%) was then mixed with steam-sterilized silversand (70%), and inoculated with 10,000 PCN eggs prior to planting a potato tuber. The un-inoculated negative controls are not included in this figure. After 13 weeks, the potato rhizosphere was collected for microbial community analysis. After 16 weeks, PCN cysts were collected and quantified. Created with BioRender.

### 2.2 Chemical analyses of soil

For each part (infected or non-infected) of each field, 2 plots were randomly selected and processed for chemical analysis. Approximately 100 g of the collected composite sample was dried overnight at 40°C. The soil samples were then crushed, and thereafter all samples were sieved using a 2 mm grid sieve. 35 g of sieved soil was used for the determination of total N, total P, organic-C, pH and % organic matter at Farm Systems Ecology (Wageningen, The Netherlands).

### 2.3 Greenhouse setup and soil sterilization

Collected (clay) soil was air-dried at ambient temperature for 2 weeks. Dried was then packed in sturdy plastic bags and crushed mechanically. One third of this crushed soil was sterilized by *γ*-irradiation (*>*30 kGy) (Synergy Health Ede B.V., The Netherlands). For each sampling point, 2L pots were filled with a mixture of 30% non-sterilized soil or sterilized soil, and 70% steam-sterilized silver sand (in quintuple). Approximately 3 weeks before the start of the experiments, potato tubers (cultivar Desiŕee, fully susceptible to *G. pallida*) were thoroughly washed with tap water to remove any adhering particles, dried with a clean paper cloth, and subsequently allowed to sprout in the dark at ambient temperature. To prepare inoculum, *G. pallida* line D383 cysts were soaked in tap water for 24 hours at room temperature and thereafter crushed using a glass rod. Egg density was determined microscopically. Before inoculation, the moisture content of each pot was brought to +/- 15%. Each pot was inoculated with 10,000 PCN eggs. Inoculation needles were used to guarantee a proper vertical inoculum distribution in each pot (Teklu et al., 2018). Nematode eggs were delivered at five positions per pot. As a positive control for PCN infectiousness, five pots filled with an artificial soil mixture optimized for PCN multiplication were inoculated with PCN. This artificial soil mixture consisted of silver sand (70% wt/wt), hydro granules (18% wt/wt, Lecaton 2 to 5 mm), and kaolin clay powder (12% wt/wt), to which NPK (12:10:18) fertilizer granules (0.001% wt/wt) were added (Teklu et al., 2018). To check for the presence of any PCN in the sampled soil, a negative control (= no inoculation with PCN) was included for all plots. After inoculation, a potato tuber containing 2 sprouts was planted in each pot. The pots were distributed in the greenhouse in a randomized complete block design. The greenhouse conditions were as follows: 20°C (day)/15 °C (night) including table cooling and +/- 70% relative humidity. The greenhouse was illuminated daily with artificial growing light from 06:30 to 09:30, after which natural sunlight provided illumination until sunset. To retain the soil moisture content around 15%, a subset of pots was weighted 3 times per week. Weight loss due to water evaporation was assessed while taking plant biomass accumulation into account, and water was added accordingly. Plants were regularly fertilized with Hyponex fertilizer (Royal Brinkman, NL).

### 2.4 Rhizosphere sampling, PCN cysts harvesting, and cyst counting

After 13 weeks, rhizosphere samples were collected from all pots, thereby avoiding co-collection of PCN cysts. Plants were carefully lifted from their pots and transversally cut plastic pipette tips (opening +/- 5 mm) were used to scrape the soil from the roots. In total, approximately 5 g of rhizosphere soil was collected per pot. The rhizosphere soil was snap frozen in liquid nitrogen immediately after collection, kept on dry ice after collection and subsequently stored at -80 °C.

16 weeks after inoculation, the cysts were harvested from all pots using the Seinhorst method (Seinhorst, 1964), cleaned with a spray nozzle over a 900 µm sieve, and thereafter air-dried at room temperature. The number of cysts and larval contents were microscopically quantified by HLB BV (Wijster, The Netherlands).

### 2.5 DNA extraction and amplification of the bacterial 16S rDNA gene

DNA was extracted from 2 g of rhizosphere soil according to Harkes et al. (2019). The DNA concentrations of all samples were determined using a Qubit (Thermo Fisher Scientific Inc.) and subsequently all DNA samples were diluted to 1 ng/µl. Nearly complete bacterial 16S rDNA was amplified using universal 27F-CM (AGAGTTTGATCMTGGCTCAG) (Frank et al., 2008) and 1492R (CGGTTACCTTGTTACGACTT) primers that were tailed with barcode sequences of the EXP-NBD196 kit (Oxford Nanopore Technologies plc., UK). PCR was performed in simplex, and each PCR reaction consisted of 12.5 µL Q5 High-Fidelity 2X MasterMix (NEB), 200 nM of each primer (IDT), 7.5 µL of UltraPure DNase/RNase-Free distilled water (Invitrogen) and 3 µL of DNA template (1 ng/µL). The following PCR scheme was used: initial denaturation at 98 °C for 30 s, followed by 25 cycles of 98 °C for 10 s, 58 °C for 30 s, 72 °C for 60 s, with a final extension step at 72 °C for 2 min. After PCR, amplification success and amplicon fragment size were verified on a 1.5% agarose gel.

### 2.6 Nanopore (16S rDNA) and Illumina (ITS2) sequencing

The 16S rDNA PCR products were merged (5 µL of each) in two pools and thereafter bead-cleaned (0.75 bead:sample ratio) using NucleoMag NGS Clean-up and Size Select beads to remove contaminants and small unwanted DNA fragments. 200 fmol of each pool was prepared for nanopore sequencing using the SQK-LSK114 kit (Oxford Nanopore Technologies plc., UK) following the manufacturer’s instructions. Libraries were then loaded on R10.4.1 flowcells (FLO-MIN114). The sequencing was performed on a MinION Mk1C and raw reads were outputted in POD5 file format.

For the characterization of the fungal community, DNA extracts were sent to Genome Qúebec (Montŕeal, Canada) for amplification of the ITS2 region (primers gITS7: GTGART-CATCGARTCTTTG and ITS4R: TCCTCCGCTTATTGATATGC), Illumina library preparation, and subsequent Illumina NextSeq (2×300bp PE) sequencing.

### 2.7 Bio-informatic analyses

The raw POD5 files resulting from the 16S nanopore sequencing were basecalled using the super-accuracy basecalling model (v5) of Dorado (v0.7.2). The resulting .fastq files were demultiplexed using Dorado and simultaneously ONT adapters and barcodes were removed. The reads were filtered on length (min. 1,000 bp, max. 2,000 bp) and quality (*>*Q20) using NanoFilt (v2.8.0; De Coster et al., 2018), prior to primer removal using cutadapt (v4.6; Martin, 2011). The taxonomic identification of the resulting reads was performed with Emu (v.3.4.5; Curry et al., 2022), a taxonomic identifier specifically developed for Nanopore reads, using the Silva 16S rDNA database (release 138.1; Quast et al., 2012). Unlike OTU or ASV-based analyses, Emu classifies individual reads and assigns them directly to a taxon, providing taxon-level read counts. The resulting Emu files were subsequently merged using the parse silva output.py Python script from the CoatOfArms GitHub repository (https://github.com/gbouras13/coatofarms). This script was modified to also output the read count data.

For the fungal communities, demultiplexed .fastq files containing raw ITS2 reads were obtained from Genome Qúebec (Montŕeal, Canada) and these reads were further processed using QIIME2 (2024.2 amplicon distribution; Bolyen et al., 2019). The fungal ITS2 region was extracted from the reads using ITSxpress (v2.0.2; Rivers et al., 2018), and in this step also the primers were removed from the reads. Next, DADA2 (v2024.2.0; Callahan et al., 2016) with no truncation was used to de-noise and merge the reads, and to remove chimeric reads. The resulting ASVs were clustered using VSEARCH (v2024.2.0; Rognes et al., 2016) as recommended for fungal ITS sequences (Kauserud, 2023; Tedersoo and Anslan, 2019). Next, the reads were taxonomically identified using the QIIME2 classify-sklearn command of the feature-classifier plugin and the pre-trained UNITE (v10.0) database (Abarenkov et al., 2024).

### 2.8 Statistical analyses

All analyses were performed in R (v4.3.1; RCoreTeam, 2013). The overall difference in cyst quantities between the putatively suppressive and conducive parts (i.e. variable ”state”) was analyzed using a Generalized Linear Model (GLM) of the MASS package (v7.3-60; Venables and Ripley, 2002). To compensate for the presence of cysts in the sampled field soil, the number of cysts in the non-inoculated negative controls of each plot was subtracted from those in the inoculated pots. Cyst quantitative data were analysed using a GLM with a Negative Biniomial (NB) distribution. For the sterilized soil data, one extreme outlier (cyst count = 44, field G, suppressive part) was removed as this prevented the use of GLM models and resulted in overdispered models. All models with a NB distribution showed no over- or underdispersion, except for the comparison of the number of living larvae for field K. For the latter analysis, a Poisson regression was used due to equidispersion of the data. Violin plots were created to visualize the cyst quantity data using ggplot2 (v3.4.2; Wickham, 2016).

The QIIME2 OTU table and taxonomy file of the fungal ITS2 reads were imported into R phyloseq objects using the qiime2R package (v0.99.6; Bisanz, 2018). Fungal OTUs with less than 10 reads were removed from the feature table. The total fungal OTU and bacterial taxa richness and mean richness for the putatively conducive and suppressive parts were determined with phyloseq (v. 1.42.0; McMurdie and Holmes, 2013). The mean richness data was checked for normality using Shapiro-Wilk tests and subsequently Student’s t-tests were used to compare the mean richness per field between the putatively conducive and suppressive parts. The bacterial mean richness data of field E was not normally distributed, and therefore a Wilcoxon rank-sum test was used in that instance. To visualize the community differences between the infected and uninfected parts of the fields, PCoA plots were created for all fields combined and each field separately using Bray-Curtis (Bray and Curtis, 1957) and robust Aitchison (Gloor et al., 2017; Martino et al., 2019) dissimilarity matrices. For the Bray-Curtis dissimilarity matrix, the OTU table was rarefied to the lowest read count (16,438 reads) using the phyloseq package. For the robust Aitchison distance, the OTU table was first rclr transformed using QsRutils (v0.1.5; Quensen, 2020) and thereafter an euclidean distance was applied when performing the ordination (Gloor et al., 2017). Subsequently, PERMANOVA tests (adonis2, vegan package v. 2.6–4; Oksanen et al., 2022) with 1,000 permutations were used to statistically determine the effect of variables block, field, and state (=putatively conducive or putatively suppressive) on the fungal communities, using both the Bray-Curtis and robust Aitchison distance. Differentially abundant taxa between the states within each field were determined by using ANCOM-BC with structural zeros detection (v1.4.0; Lin and Peddada, 2020) and DESeq2 (v1.40.2; Love et al., 2014) with the ’poscount’ size factor estimator to account for features with zero counts. The significance threshold of the Holm (ANCOMBC) and Benjimani-Hochberg (DESeq2) adjusted P-values was set at 0.05. For improved visualization, the DESeq2 log2 fold changes where shrank using the the ’apeglm’ estimator. The differential abundance results were visualized by creating a heatmap using the ComplexHeatmap package (v.2.16.0; Gu, 2022). Only putatively fungal taxa that are known to harbor nematode antagonists are shown. The taxa table resulting from the bacterial 16S rDNA sequencing was analyzed using the same methods as those applied to the fungal ITS2 OTU table, with the following differences: a minimum abundance threshold of 0.0001 was used to retain taxa, and a rarefaction depth of 110,000 reads was used for Bray-Curtis dissimilarity.

For the identification of putatively nematode antagonists, we listed bacterial and fungal genera that were included in three reviews, namely Topalovíc et al. (2020), Li et al. (2015), and Stirling (2014). It is noted that some genera are dominated by nematode antagonists (*e.g*., the bacterial genus *Pasteuria*), while for other genera nematode antagonists constitute a small minority only (*e.g.,* the bacterial genus *Pseudomonas*). This list includes genera with representatives that directly parasitize plant-parasitic nematodes (PPNs), as well as microbial taxa that are known to promote PAMP-triggered immunity in their host plant

## 3 Results

### 3.1 Verification of PCN suppressiveness under controlled conditions

To see whether apparent PCN suppressiveness under field conditions could be reproduced under controlled greenhouse conditions, PCN multiplication on sterilized and non-sterilized soil from putatively conducive and suppressive patches was compared. The overall number of cysts in the non-sterilized soils, quantified at the end of the experiment, was 21% lower (GLM.NB, P=0.042) in the putatively suppressive parts of the field than in the conducive parts (Figure 2A). A subsequent per field comparison of PCN multiplication showed that for field E (GLM.NB, P=0.025) and Field K (GLM.NB, P=0.003) the number of cysts quantified in the putatively suppressive part was respectively 35% and 34% lower than in the conducive part (Figure 2B). For field G (GLM.NB, P=0.17) and field S (GLM.NB, P=0.53), there were no significant differences in the average number of cysts between the field parts. When the soil was *γ*-sterilized, the observed overall difference in PCN multiplication between the putatively conducive and suppressive soil was no longer present (GLM.NB, P=0.42) (Figure 2C). Field K (GLM.NB, P=0.097) showed no differences in cyst quantities between putatively suppressive and conducive parts anymore after *γ*-sterilization, and in field E the reduction in PCN multiplication in the putatively suppressive soil was also nullified (Figure 2D). Therefore, we conclude that the observed suppressiveness was caused by biota present in these soils. For field S, the number of cysts in the sterilized putatively suppressive soil was 14% lower (GLM.NB, P=0.03) as compared to soil from the putatively conducive field part. This suggests that heterogeneous edaphic factors in Field S might have contributed to the observed reduction in PCN manifestation.

**Figure 2:**
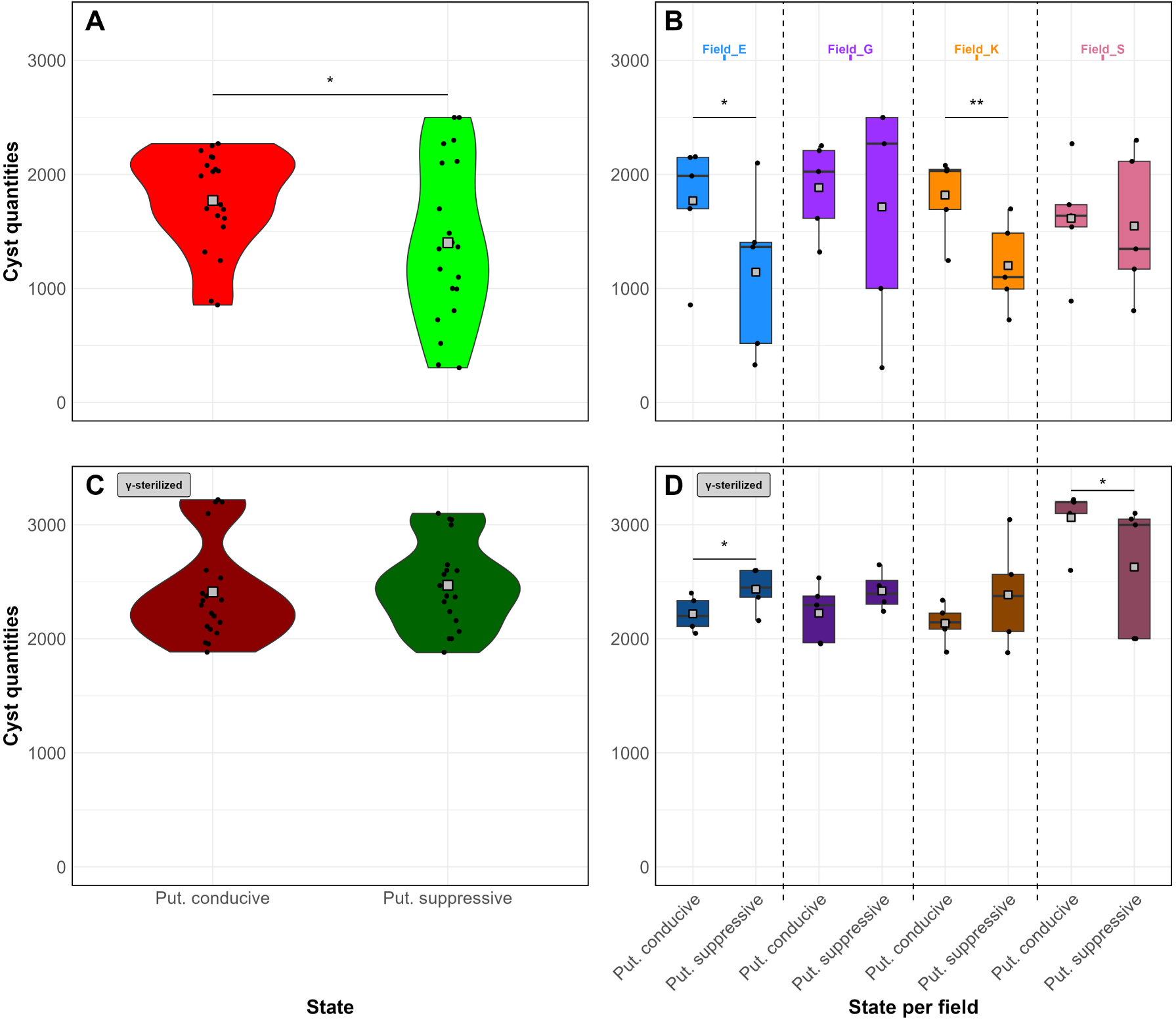
Number of potato cyst nematode (PCN) cysts per pot quantified at 16 weeks post-inoculation on fully susceptible potato plants. A) Cyst counts upon exposure to soil from putatively suppressive (green) and conducive (red) field patches from four different locations together (n=20) and B) PCN multiplication per field (n=5). C) and D) PCN multiplication upon exposure to gamma-sterilized putatively conducive (dark red) and suppressive (dark green) soil across all fields (n=20) (C) and per field (n=5) (D). * (P *<*0.05) and ** (P *<*0.01) indicate statistical significant difference as determined by a GLM with a Negative Binomial distribution. Put. = putatively.

Sterilization of the putatively suppressive soil increased PCN multiplication in fields E (113%, GLM.NB, P*<*0.00001), K (99%, GLM.NB, P*<*0.00001), and S (70%, GLM.NB, P=0.0003) (Supplementary Figure S1-B). For field G, sterilization increased the number of PCN cysts by 41%, but this change was not significant (GLM.NB, P=0.075). Although to a lesser extent, sterilization also increased the number of PCN cysts on potato plants exposed to conducive soil (Supplementary Figure S1-D). Thus, *γ*-sterilization had a larger impact on PCN multiplication in soil from putatively suppressive field parts than in soil from conducive field parts.

Next to the number of cysts, the overall number of living larvae per cyst was counted. This parameter did not significantly differ between soil from putatively conducive and suppressive patches (GLM.NB, P=0.58, Supplementary Figure S2-A). Field K was exceptional as cysts from pots with putatively suppressive soil comprised 8.5% (GLM.NB, P=0.049) less living larvae as compared to conducive soil (Supplementary Figure S2-B).

### 3.2 Conducive and suppressive field sections show contrasting fungal and bacterial communities

The complete bacterial 16S rDNA and fungal ITS2 region were sequenced to analyse the microbial rhizosphere communities (mean reads per sample respectively 167,455 and 79,191). After filtering, we identified a total of 1,431 bacterial and 560 fungal taxa for all fields (Table 1). Per field, a total of 174-341 fungal and 865-1,193 bacterial taxa were identified (Table 1). No significant differences in mean fungal OTU richness were observed between the putatively conducive and suppressive field parts. The mean bacterial richness was higher for the putatively suppressive field parts when the data from the four locations were aggregated (10.4% increase, Student’s t-test, P= 0.016). With a richness increase of 39.2% (Wilcoxon rank-sum test, P=0.009) as compared to the putatively conducive field part, field E was exceptional.

**Table 1:** Number of fungal OTUs and bacterial taxa (see 2.7 Bio-informatic analyses) after filtering low abundant taxa. Total number of OTUs and taxa are given for all fields aggregated and for each field separately. Mean OTU and taxa numbers are presented for the putatively conducive and suppressive field sections. Letters are used to indicate significant differences between the mean OTUs and taxa per field (i.e., row-wise per taxonomic group), according to the student t-test (P *<* 0.05)

|  | Fungal OTUs |  |  | Bacterial taxa |  |  |
| --- | --- | --- | --- | --- | --- | --- |
|  | Total # of OTUs | Mean # of OTUs |  | Total # of taxa | Mean # of taxa |  |
|  |  | Put. con | Put. sup |  | Put. con | Put. sup |
| Fields aggregated | 560 | 57<sup>a</sup> | 65<sup>a</sup> | 1431 | 550<sup>a</sup> | 607<sup>b</sup> |
| Field E | 341 | 78<sup>a</sup> | 93<sup>a</sup> | 1193 | 483<sup>a</sup> | 672<sup>b</sup> |
| Field G | 207 | 42<sup>a</sup> | 69<sup>a</sup> | 874 | 568<sup>a</sup> | 591<sup>a</sup> |
| Field K | 174 | 50<sup>a</sup> | 37<sup>a</sup> | 875 | 592<sup>a</sup> | 593,6<sup>a</sup> |
| Field S | 215 | 60<sup>a</sup> | 62<sup>a</sup> | 865 | 557<sup>a</sup> | 571<sup>a</sup> |

PCoA plots (Bray-Curtis dissimilarity) were generated to visualize differences in the overall fungal (Figure 3) and bacterial (Figure 4) communities between the putatively conducive and putatively suppressive field parts (*i.e.,* variable ’state’) of all fields under investigation In general, the fungal communities of each of the four fields under unvestigation clustered together and PERMANOVA indicated a significant effect of field on the composition of the rhizobiome (P*<*0.001, R^2^ = 0.20) (Figure 3A). Field-specific community comparisons revealed the presence of distinct fungal communities in the putatively conducive and suppressive parts of each field (see PERMANOVA results, Figure 3B-D). For all fields, the variable ”state” (*i.e.* putatively conducive or suppressive) explained a substantial part of the variation in fungal communities (R^2^*>*20%).

**Figure 3:**
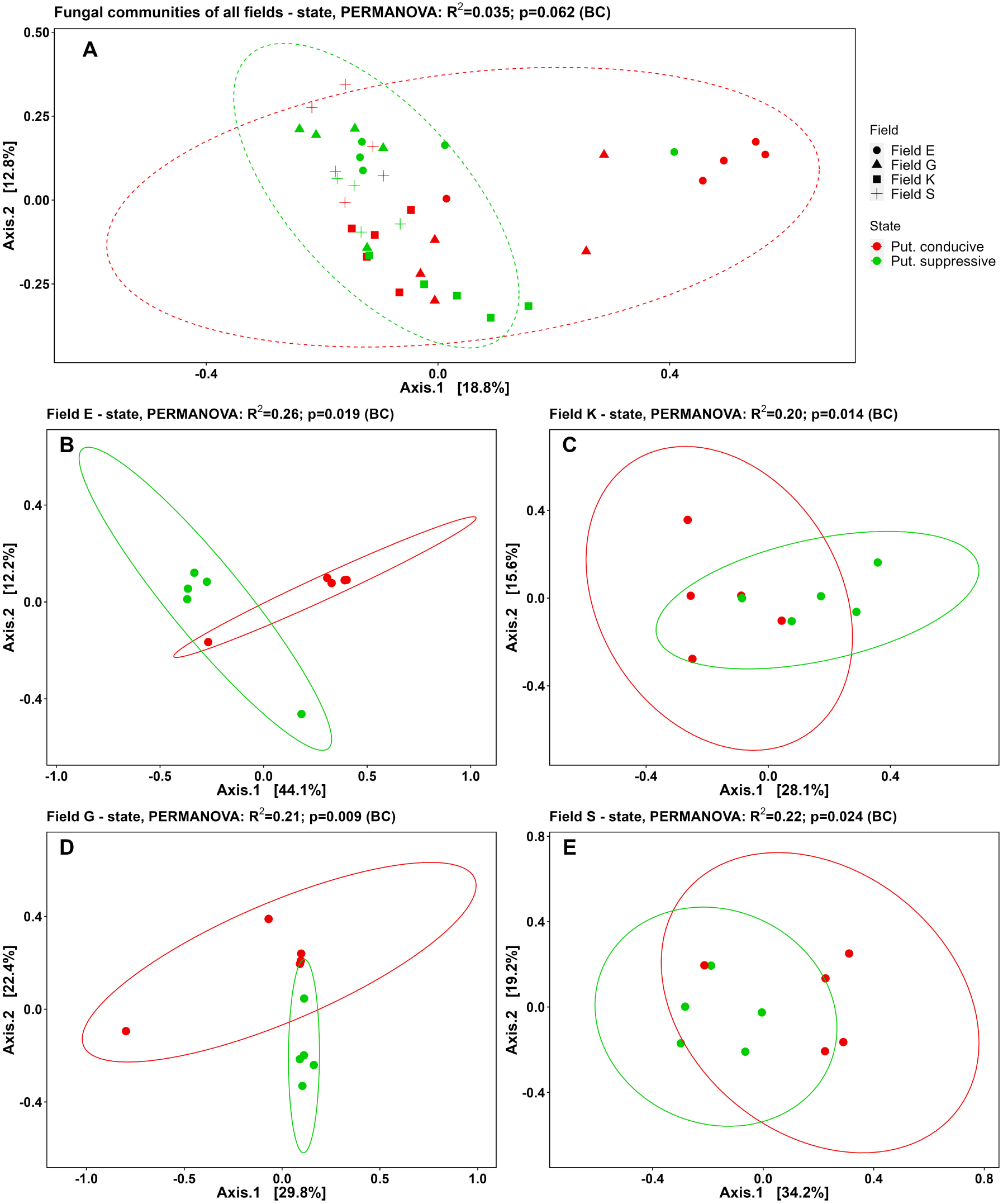
PCoA plots based on the Bray-Curtis dissimilarity matrix of fungal communities in rhizosphere samples collected from potato plants 13 weeks after inoculation with potato cyst nematodes. A) Fungal communities in rhizosphere of plants exposed to putatively conducive (red) or suppressive (green) soil from all four field locations and B-E) at individual field location level (n=5). B = field E, C = field K, D = field G, and E = field S. Next to A), symbols for the individual field locati3o5ns are presented. Figure headers include the PERMANOVA result of factor ’state’ (= putatively suppressive or conducive). Put. = putatively.

**Figure 4:**
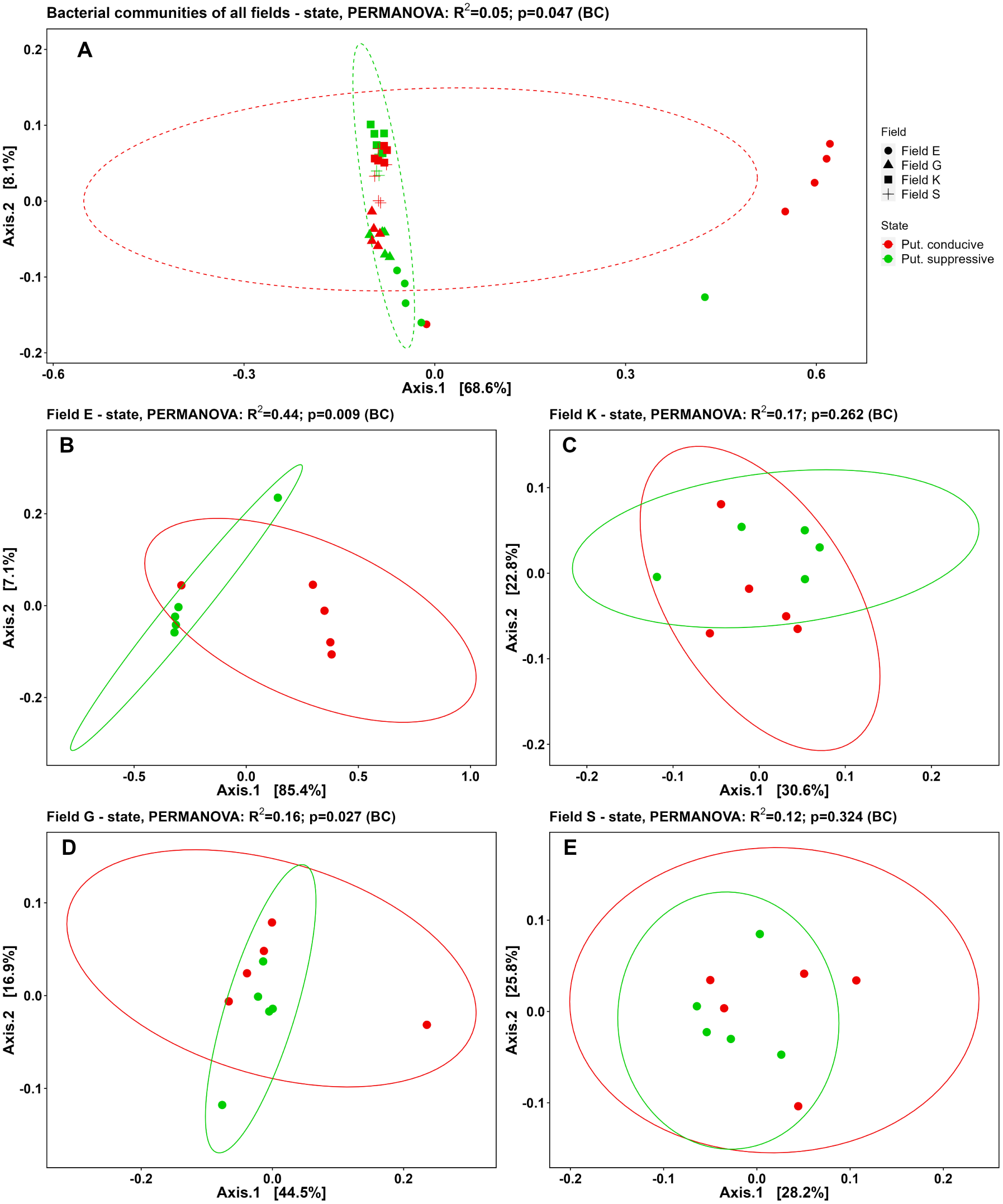
PCoA plots based on the Bray-Curtis dissimilarity matrix of the bacterial communities in rhizosphere samples collected from potato plants 13 weeks after inoculation with potato cyst nematodes. A) Bacterial communities in rhizosphere of plants exposed to putatively conducive (red) or suppressive (green) soil from all four field locations, and B-E) at individual field location level (n=5). B = field E, C = field K, D = field G, and E = field S. Next to A), symbols for the individua3l6field locations are presented. Figure headers include the PERMANOVA result of factor state (= putatively suppressive or conducive). Put. = putatively.

Also the composition of the bacterial communities was significantly different between the four field locations (P*<*0.001, PERMANOVA) (Figure 4A). Analysis of individual fields showed a clear separation in community composition between putatively suppressive and conducive soil in fields E and G (Figures 4B and D, respectively) (PERMANOVA, P*<*0.05). The bacterial community composition in fields K and S (Figures 4C and E, respectively ) was not significantly affected by its state. PCoA plots and PERMANOVA analyses were also generated using the Aitchison distance and this produced similar results for the fungal and bacterial communities (respectively Supplementary Figures S3 and S4) except for the fungal communities in fields K and S. These results show that the fungal and bacterial rhizosphere communities from potato plants grown in putatively suppressive and conducive soil for 13 weeks differ in composition.

### 3.3 Intra- and interfield composition of nematode antagonist communities

Two distinct algorithms, ANCOM-BC and DESeq2, were used to determine which fungal and bacterial taxa are differentially abundant in the suppressive and conducive part of each field. When data from all field locations were aggregated, ANCOM-BC revealed 1 fungal taxon and DESeq2 17 fungal taxa that were differentially abundant between the two distinct states. The latter 17 taxa included the nematode antagonists *Arthrobotrys elegans* (more abundant in putatively suppressive state) and *Trichoderma hamatum* (less abundant in putatively suppressive state). Differential abundance analyses at field level revealed a higher abundance of fungal taxa known to interact with plants in some of the putatively suppressive field sections. For example, the arbuscular mycorrhizal fungus *Claroideoglomus claroideum* and the endophyte *Coniochaeta sp.* were more abundant in the putatively suppressive part of field E (Supplementary Figure S5). Various fungal nematode antagonists were differentially abundant in the putatively suppressive parts of the fields (Figure 5). ANCOM-BC (Figure 5A) analysis revealed that in field E, *Arthrobotrys elegans* (syn: *Orbilia elegans*, *A. oudemansii* ; Scholler et al., 2000; Baral et al., 2020), *Acremonium spp*, and an Orbiliaceae species were more abundant in the putatively suppressive part. In field G, *Arthrobotrys elegans* was also more abundant. In field K, *Metacordyceps chlamydosporia* (synonymous to *Pochonia chlamydosporia*) was more abundant, while an increase in presence of *Purpureocilium lilacinum* was observed in field S. These results were corroborated by DESeq2 (Figure 5B). Additionally, DESeq2 identified several other nematode antagonists in each field that were more abundant in the putatively suppressive parts of the fields, *e.g., Metapochonia suchlasporia* (field G & K) and *Trichoderma harzianum* (Field S). Both analyses also identified fungal antagonists that were less abundant in the putatively suppressive parts in each field, *e.g., T. hamatum* (field E) and *A. conoides* (field G). Furthermore, some of the the nematode antagonists showed inter-field differences in differential abundance between the two states. For example, *P. lilacinum* was more abundant in the putatively suppressive parts of field K and S, while this fungi was less abundant in this part of field E (DESeq2). These results show that various fungal nematode antagonists are associated with the putatively suppressive parts of each field and suggest that this is field specific.

**Figure 5:**
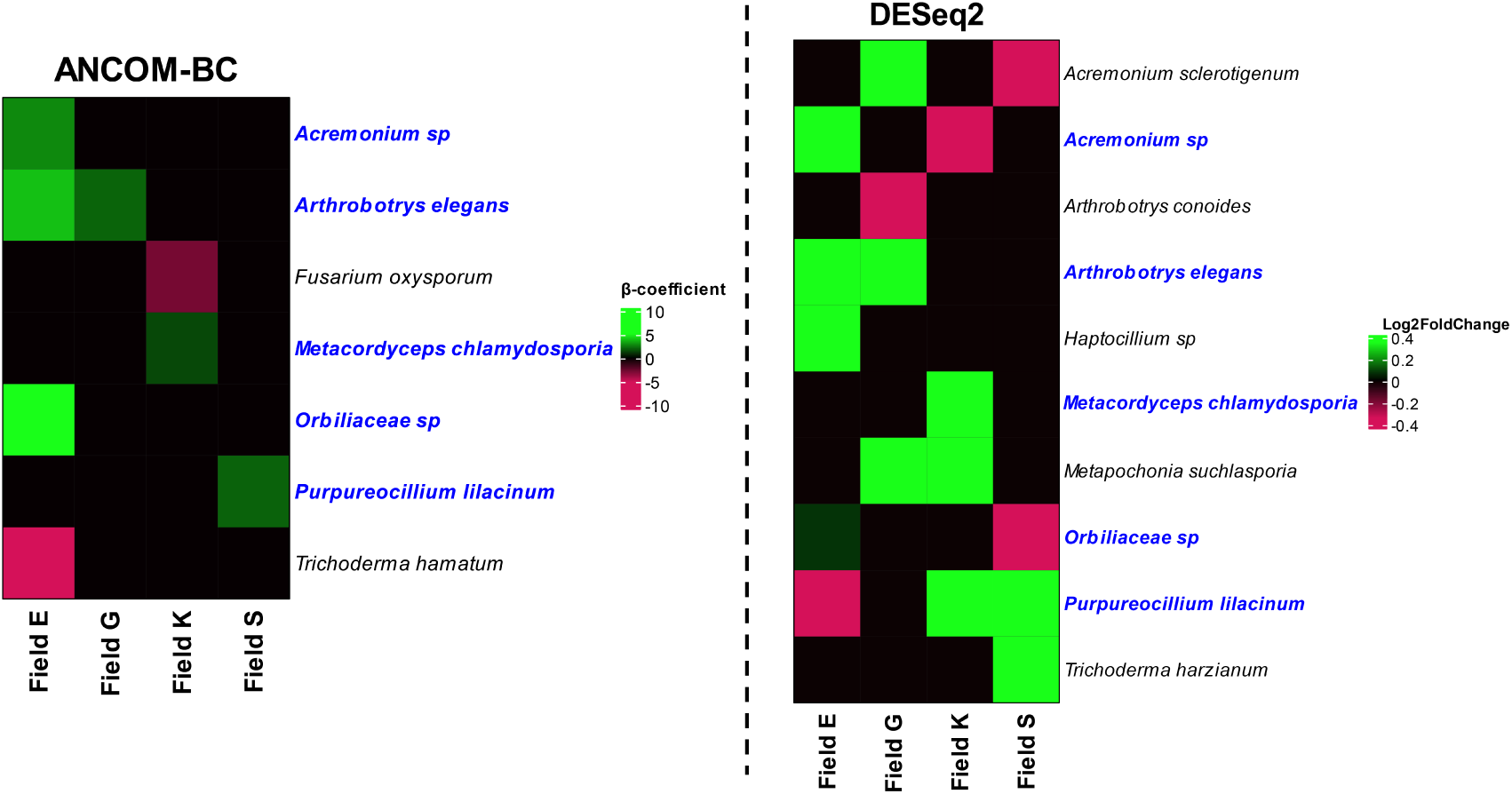
Differentially abundant (putative) fungal nematode antagonists as revealed by ANCOM-BC (left) or DESeq2 (right) (adjusted P-value *<*0.05). Taxa with a positive β-coefficient or log2 fold change (= green) are more abundant in the putatively suppressive field parts, while taxa with a negative β-coefficient or log2 fold change (= red) are less abundant in the putatively suppressive field part. Taxa indicated in blue were differentially abundant in both analyses.

When data from all field locations were aggregated, ANCOM-BC revealed 6 and DESeq2 10 bacterial taxa that were differentially abundant between the two distinct states. These differentially abundant taxa did not include known nematode antagonists. At field level, ANCOM-BC analyses identified differentially abundant bacterial taxa (Figure 6) that are known as nematode antagonists and representives of bacterial genera that are known to comprise antagonists (e.g., Topalovíc et al., 2020; Li et al., 2015; Stirling, 2014). Although these bacterial genera comprise nematode antagonists, no information is available about the trophic ecology of some of these species and therefore they are labeled as ”potential nematode antagonists”. Nearly complete 16S rRNA sequences allowed us to identify nearly all potential bacterial antagonists at species-level. Two of the differentially abundant bacterial species are known nematode antagonists *Pseudomonas fluorescens* and *P. putida*, and these species were more abundant in the putatively suppressive parts of field E. The largest other contrasts were found in field E, where several species of *Lysobacter*, and *Streptomyces* were more abundant in the putatively suppressive part, while an *Arthrobacter* and *Bacillus* species were under represented. In field G and field K two potential nematode antagonists, respectively *Pseudomonas migulae* and *Streptomyces polymachus*, were less abundant in the putatively suppressive field parts. In field S, *Bacillus murimartini* was shown to be more abundant in the putatively suppressive part of the field.

**Figure 6:**
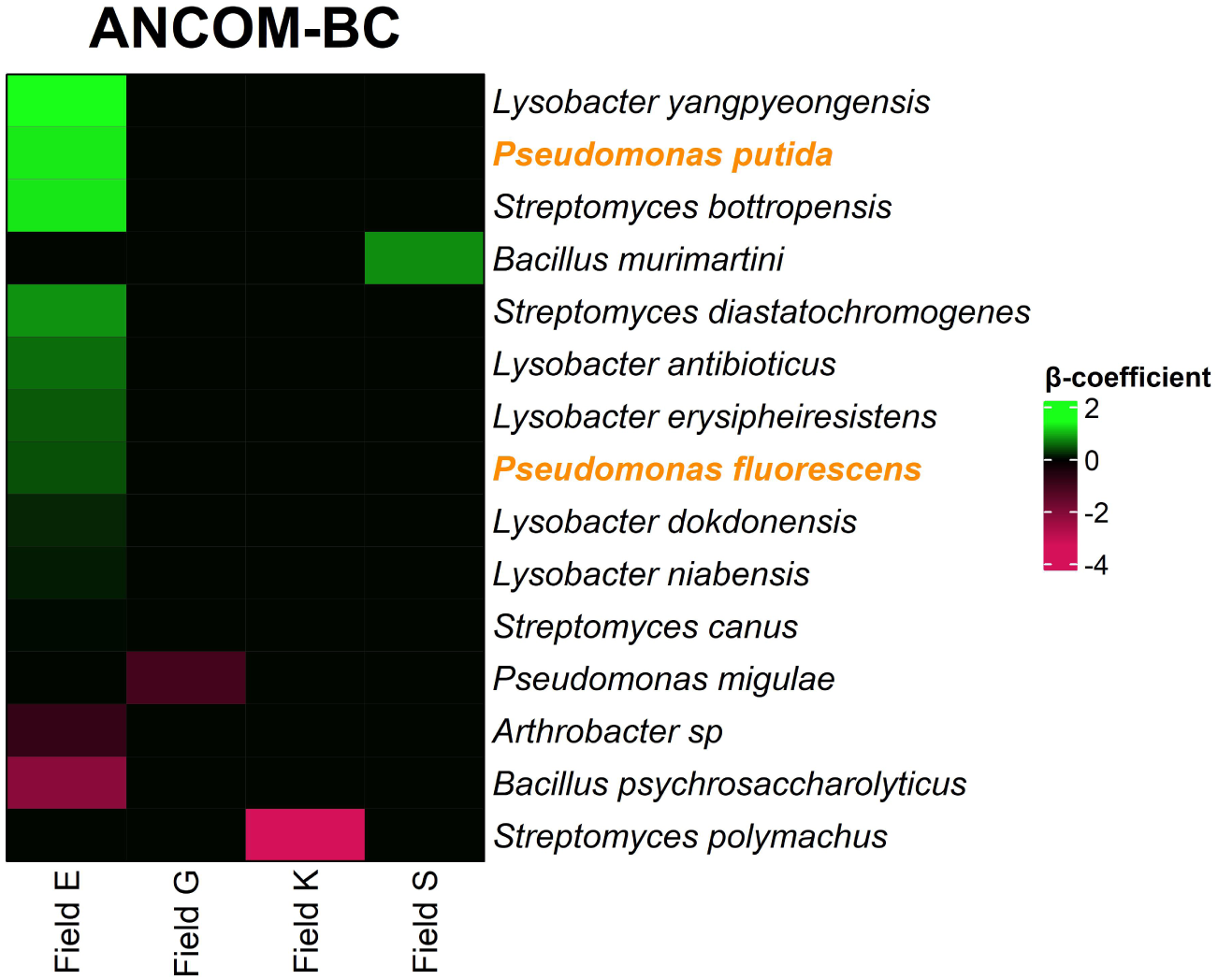
Differentially abundant putative and potential bacterial nematode antagonists as revealed by ANCOM-BC (Holm adjusted P-value *<*0.05). Bacterial species for which nematode-antagonistic strains have been described are highlighted in orange. The other taxa (black) represent bacterial genera that harbor nematode antagonists, whereas no relevant information is available for the specific species. Such taxa are labeled as ”potential antagonists”. Taxa with a positive β-coefficient (= green) are more abundant in the putatively suppressive field parts, while taxa with a negative β-coefficient fold change (= red) are less abundant in the putatively suppressive field parts.

### 3.4 Abiotic description of the fields

Chemical analysis of 2 samples of each state of each field revealed no obvious differences in total N, total P, organic matter, organic carbon, and moisture (Supplementary Figure S6). Notably, a pH difference was observed between the states of field E.

## 4 Discussion

Natural suppression of soil-borne plant pathogens by microbiota is an interesting and biologically complex phenomenon. Comparison of microbial communities from conducive and pathogen-suppressive fields as a means to pinpoint this phenomenon seems straightforward, but relevant biological signals could be obscured by variability in field conditions unrelated to disease suppressiveness. Due to their quarantine status, potato cyst nematodes (PCN) are extensively monitored in potato production areas and, depending on local phytosanitary legislation, within-field PCN distribution data might be available. Among distribution data from numerous fields, we selected four fields that showed patchy PCN distributions that couldn’t be attributed to obvious local field variables. Upon testing PCN reproduction on soils from putatively conducive and suppressive field patches under greenhouse conditions, local PCN suppressiveness as well as its biological origin were confirmed. Characterization of microbial rhizosphere communities revealed nematode antagonists positively associated with this local PCN suppression. Notably, the conglomerate of antagonists associated with PCN suppression appeared to be specific for each of the fields.

### 4.1 The biotic origin of local PCN suppression

*γ*-irradiation-based inactivation of the soil microbial community was used to verify the biotic origin of PCN suppression. This verification approach has previously also been used to show biotic of suppression of the fungus *Fusarium culmorum* on wheat (Ossowicki et al., 2020), and of the root-knot nematode *Meloidogyne javanica* on cucumber (Watson et al., 2020). Comparison of the sterilized and non-sterilized soil of the conducive state, also showed an increase of the number of PCN cysts in the conducive soil upon sterilization. However, the percentage of increase and effect size as determined by the ”estimate” was substantially lower as compared to the suppressive soils. We hypothesize that sterilization of the soil eliminated any biological competitors of the inoculated PCN. Thus, the dominant microbial community in the untreated conducive soil contributed to a mild general suppression effect (Weller et al., 2022).

Suppressiveness could result from a reduced number of cysts formed, but also from a reduction in the number of viable content per cyst (Eberlein et al., 2016; Indarti et al., 2021; Nagachandrabose, 2020). In our study, the overall number of cysts was reduced in the putatively suppressive soil, but the overall viable content of the cysts (*i.e*., number of living larvae) was not significantly affected. This suggests that due to biotic factors, fewer juveniles could reach and/or successfully penetrate the roots of a suitable host (*i.e*., fewer cysts). Once infective J-2’s had established a feeding site, the soil microbiome from PCN suppressive field appeared to have no effect on fertilization and subsequent maturation of fertilized eggs.

### 4.2 Nematode antagonists associated with suppressive field patches

Characterization and comparison of bacterial and fungal communities in putatively conducive and suppressive parts of the fields revealed various fungal and bacterial nematode-antagonistic taxa that were more abundant in the putatively suppressive parts. Outside of the center of PCN diversity (Andean region in South America), PCN suppressive soils are thought to be uncommon (Kerry et al., 2009). In contrast to *Meloidogyne* suppressive soils (*e.g.,* Watson et al., 2020; Adam et al., 2014; Bell et al., 2016), little is known about local soil microbiomes that could suppress PCN. Nevertheless, multiple bacterial and fungal isolates from soils originating from Ireland (Cronin et al., 1997), Portugal (Santos et al., 2013), Tunisia (Hajji et al., 2017), and India (Nagachandrabose, 2020) have been identified as PCN antagonists. Furthermore, a recent survey identified several (abundant) taxa in a PCN suppressive soil from Scotland (UK) (PhD thesis Jones, 2024), including those that were abundant in the putatively suppressive field sections in our study (*e.g., Arthrobotrys sp.* and *Streptomyces sp.*).

Several known PPN antagonists were more abundant in the putatively suppressive soils included in our study, such as *Purpureocillium lilacinum*, *Arthrobotrys elegans*, *Metacordyceps chlamydosporia*, and *Pseudomonas fluorescence*. The potential of the nematode-egg parasite *P. lilacinum* for the control of PPNs - including PCN - has been well documented (*e.g.,* Hajji et al., 2017; Singh et al., 2013; Nagachandrabose, 2020). Accordingly, several microbial bio-formulations containing *P. lilacinum* are used as bionematicides in PCN management (Mhatre et al., 2022). The capacity to form nematode-trapping adhesive networks is conserved across the genus *Arthrobotrys* (Scholler et al., 1999) and numerous members of *Arthrobotrys* have been identified to trap mobile juveniles of various PPNs (*e.g., M. incognita*; Soliman et al., 2021; *Pratylenchus zeae*; Sankaranarayanan and Hari, 2021; *Xiphnema index* ; Wernet and Fischer, 2023). *Metacordyceps chlamydosporia* parasitizes PCN eggs (Manzanilla-Ĺopez et al., 2017) and *Pseudomonas fluorescence* is able to reduce PPN juvenile mobility (Cronin et al., 1997). Hence, these fungal and bacterial taxa collectively have the capacity to parasitize PCN eggs and reduce the number of juveniles that reach the roots, resulting in a reduced number of cysts. Apparently, these microbes did not interfere with the induction or maintenance of PCN induced feeding structures (syncytia) as this would have affected the viable content of the cysts. Also several fungal nematode antagonists were more abundant in the putatively suppressive patches of fields were PCN multiplication was not reduced in the greenhouse assay. The lifestyle (*i.e*., saprotrophic or nematophagous) that many fungal nematode antagonists exert strongly depends on the environmental conditions (*e.g.,* Nordbring-Hertz, 2004). Thus, the suppressive potential of these soils may decrease under altered environmental conditions in the pots.

Similarly to PCN, the soybean cyst nematode *Heterodera glycines*, has spread from its center of origin (north-east China) to nearly all major soybean producing regions in the world (Yu, 2011). Also for soybean - soybean cyst nematode disease suppressiveness has been documented outside the center of diversity. For example, Chen (2007) and Hu et al. (2017) identified *H. glycines* suppressive soils in the United States and identified bacterial (*e.g., Lysobacter spp.*) and fungal (e.g., *Hirsutella rhossiliensis*) taxa associated with this suppressive capacity. These studies corroborate our findings that local microbial communities might be able to suppress non-native soil-borne pathogens.

Not all nematode-antagonistic taxa present in putatively suppressive field patches were stimulated upon controlled exposure to PCN-infected potato plants. Whereas *Pseudomonas sp.* and *Acremonium sp.* were more abundant in the putatively suppressive part of field E, the abundance of *Trichoderma hamatum*, an antagonistic fungus able to parasitize PCN eggs (Nyang’au et al., 2023) and *Arthrobacter spp.*, a bacterial genus secreting nematode antagonistic volatile organic compounds (Topalovíc et al., 2020), were suppressed. These results possibly point at competition between PPN antagonists (Thakur and Geisen, 2019) and/or differences in feeding strategy of antagonists.

### 4.3 Dissimilarity of antagonistic microbial assemblages between geographically close fields

Although we acknowledge that the number of fields investigated here is limited, we observed that the composition of the microbial community associated with PCN suppression differed substantially among fields. Contrasts in soil microbiome composition between different agricultural fields in the same geographical area are common (*e.g.,* Castillo et al., 2017), including taxa associated with fields showing distinct levels of (nematode) suppressiveness (Watson et al., 2020; Zhou et al., 2019; Ossowicki et al., 2020). For example, Watson et al. (2020) found a positive correlation of the bacterial order Chthoniobacterales with a particular *Meloidoigyne* suppressive site in Florida (USA), while this same taxon was suppressed in another nearby suppressive site.

The soil microbial community composition is mainly driven by abiotic factors such as pH (Rousk et al., 2010) and organic matter composition (Sun et al., 2016). Therefore, even subtle variation in soil types, soil chemical properties, and soil management (Rousk et al., 2010; Lauber et al., 2008; Harkes et al., 2019) across fields might have co-shaped the set of antagonists observed in our fields. Apart from soil physical and chemical properties, also ecological processes such as top-down regulation by microbial predators (Thakur and Geisen, 2019) and stochastic soil processes (*e.g.,* priority effects, Powell et al., 2015; Benucci et al., 2023) might have contributed to the shaping of these distinct PCN antagonistic microbial conglomerates in geographically close fields.

### 4.4 Conclusion

Local within-field variation in disease suppression could be instrumental in pinpointing the microbial basis of disease suppressive soils. In this respect, the quarantine status of PCN was advantageous in our study, as statutory monitoring obligations resulted in the availability of high-resolution distribution data. Our findings highlight that the local differences in soil microbiome composition can influence the local establishment of a phytopathogen within a field. While we show the impact of local variation in the soil microbiome, further research should pinpoint the cause of this localized disease suppression. Such insights will advance our understanding of the manipulability and controllability of disease suppressive soils. Distinct conglomerates of antagonists were associated with PCN suppression even in geographically close fields. This finding emphasizes the challenges associated with the application of biological control agents to manage PCN and other soil-borne pathogens. Finally, confirmation of the occurrence of PCN suppressive soils in Europe encourages further efforts to exploit this potential as a new strategy to durably manage this notorious plant pathogen.

## Data availability statement

Raw demultiplexed 16S nanopore reads and ITS2 Illumina reads are openly accessible in the Sequence Read Archive (SRA) of NCBI under BioProject accession PR-JNA1217848.

## Acknowledgements

We thank the 3 anonymous farmers very much to allow us to sample their fields and their willingness to share field information. We would like to thank Misghina Goitom Teklu (WUR) for the valuable discussions regarding PCN; Luc van Pierre (WUR) for the assistance with field sampling and inoculation of the pots; Unifarm (WUR) for their assistance with crushing the soil; Gijs de Wildt (WUR) for his assistance with rhizosphere sampling; Max Jenje, Axel van der Vooren, and Thomas Moolenaar (WUR) for their assistance with cyst extraction; The Q-organisms team of HLB for their assistance with cyst quantification. The graphical abstract was created with BioRender.

## Author contributions

**R. van Himbeeck:** Formal analysis, Investigation, Methodology, Visualization, Writing - original draft, Writing - review & editing. **S. Geisen:** Funding acquisition, Conceptualization, Methodology, Writing - review & editing. **C. van Schaik:** Investigation, Writing - review & editing. **S. van den Elsen:** Investigation, Writing - review & editing. **R. Berendsen:** Investigation, Writing - review & editing. **A. Bertran:** Investigation, Resources, Writing - review & editing. **E. Schepel:** Investigation, Resources, Writing - review & editing. **J. Helder:** Conceptualization, Funding acquisition, Methodology, Supervision, Writing - review & editing.

## References

Abarenkov, K. et al. (2024). The UNITE Database for Molecular Identification and Taxonomic Communication of Fungi and Other Eukaryotes: Sequences, Taxa and Classifications Reconsidered. Nucleic Acids Research 52.D1, pp. D791–D797. issn: 0305-1048. doi: 10.1093/nar/gkad1039.

Abo-Elyousr, K. A. M. and Hassan, S. A. (2021). Biological Control of Ralstonia Solanacearum (Smith), the Causal Pathogen of Bacterial Wilt Disease by Using Pantoea Spp. Egyptian Journal of Biological Pest Control 31.1, p. 113. issn: 2536-9342. doi: 10.1186/s41938-021-00460-z.

Adam, M., Westphal, A., Hallmann, J., and Heuer, H. (2014). Specific Microbial Attachment to Root Knot Nematodes in Suppressive Soil. Applied and Environmental Microbiology 80.9. Ed. by J. L. Schottel, pp. 2679–2686. issn: 0099-2240, 1098-5336. doi: 10.1128/AEM.03905-13.

Baral, H.-O., Weber, E., and Marson, G. (2020). Monograph of Orbiliomycetes (Ascomycota) Based on Vital Taxonomy: Part I + II. Luxembourg: National Museum of Natural History Luxembourg. isbn: 978-2-919877-24-9.

Bell, N. L. et al. (2016). Detection of Invertebrate Suppressive Soils, and Identification of a Possible Biological Control Agent for Meloidogyne Nematodes Using High Resolution Rhizosphere Microbial Community Analysis. Frontiers in Plant Science 7. issn: 1664-462X. doi: 10.3389/fpls.2016.01946.

Benucci, G. M. N., Beschoren da Costa, P., Wang, X., and Bonito, G. (2023). Stochastic and Deterministic Processes Shape Bioenergy Crop Microbiomes along a Vertical Soil Niche. Environmental Microbiology 25.2, pp. 352–366. issn: 1462-2920. doi: 10.1111/1462-2920.16269.

Bisanz, J. E. (2018). qiime2R: Importing QIIME2 Artifacts and Associated Data into R Sessions. Version 0.99 13.

Bolyen, E., Rideout, J. R., Dillon, M. R., Bokulich, N. A., Abnet, C. C., Al-Ghalith, G. A., Alexander, H., Alm, E. J., Arumugam, M., and Asnicar, F. (2019). Reproducible, Interactive, Scalable and Extensible Microbiome Data Science Using QIIME 2. Nature biotechnology 37.8, pp. 852–857.

Bray, J. R. and Curtis, J. T. (1957). An Ordination of the Upland Forest Communities of Southern Wisconsin. Ecological Monographs 27.4, pp. 326–349. issn: 0012-9615. doi: 10.2307/1942268.JSTOR:1942268.

Bubici, G., Kaushal, M., Prigigallo, M. I., Gómez-Lama Cabańas, C., and Mercado-Blanco, J. (2019). Biological Control Agents Against Fusarium Wilt of Banana. Frontiers in Microbiology 10. issn: 1664-302X. doi: 10.3389/fmicb.2019.00616.

Callahan, B. J., McMurdie, P. J., Rosen, M. J., Han, A. W., Johnson, A. J. A., and Holmes, S. P. (2016). DADA2: High-resolution Sample Inference from Illumina Amplicon Data. Nature Methods 13.7, pp. 581–583. issn: 1548-7105. doi: 10.1038/nmeth.3869.

Castillo, J. D., Vivanco, J. M., and Manter, D. K. (2017). Bacterial Microbiome and Nematode Occurrence in Different Potato Agricultural Soils. Microbial Ecology 74.4, pp. 888– 900. issn: 1432-184X. doi: 10.1007/s00248-017-0990-2.

Cazzaniga, S. G., Belliard, P., van Steenbrugge, J., Van den Elsen, S., Lombaers, C., Visser, J., Molendijk, L., Macia-Vicente, J. G., Postma, J., and Mommer, L. (2025). On the Diversity of Nematode Antagonists in an Agricultural Soil, and Their Steerability by Root-Knot Nematode Density and Cover Crops. Soil Biology and Biochemistry 202, p. 109693.

Cazzaniga, S. G., Braat, L., Elsen, S. van den, Lombaers, C., Visser, J., Obinu, L., Macía-Vicente, J. G., Postma, J., Mommer, L., and Helder, J. (2023). Pinpointing the Distinctive Impacts of Ten Cover Crop Species on the Resident and Active Fractions of the Soil Microbiome. Applied Soil Ecology 190, p. 105012.

Chen, S. (2007). Suppression of Heterodera Glycines in Soils from Fields with Long-Term Soybean Monoculture. Biocontrol Science and Technology. doi: 10.1080/09583150600937121.

Cronin, D., Moenne-Loccoz, Y., Fenton, A., Dunne, C., Dowling, D. N., and O’gara, F. (1997). Role of 2,4-Diacetylphloroglucinol in the Interactions of the Biocontrol Pseudomonad Strain F113 with the Potato Cyst Nematode Globodera Rostochiensis. Applied and Environmental Microbiology 63.4, pp. 1357–1361. doi: 10.1128/aem.63.4.1357-1361.1997.

Curry, K. D. et al. (2022). Emu: Species-Level Microbial Community Profiling of Full-Length 16S rRNA Oxford Nanopore Sequencing Data. Nature Methods 19.7, pp. 845–853. issn: 1548-7105. doi: 10.1038/s41592-022-01520-4.

De Coster, W., D’Hert, S., Schultz, D. T., Cruts, M., and Van Broeckhoven, C. (2018). NanoPack: Visualizing and Processing Long-Read Sequencing Data. Bioinformatics 34.15, pp. 2666–2669. issn: 1367-4803. doi: 10.1093/bioinformatics/bty149.

Eberlein, C., Heuer, H., Vidal, S., and Westphal, A. (2016). POPULATION DYNAMICS OF GLOBODERA PALLIDA UNDER POTATO MONOCULTURE. Nematropica 46.2, pp. 114–120. issn: 2220-5616.

European Union (2022). EUR-Lex - 32022L0185 - EN - EUR-Lex. url: https://eur-lex.europa.eu/legal-content/EN/TXT/?uri=uriserv%3AOJ.L_.2022.185.01.0012.01.ENG&toc=OJ%3AL%3A2022%3A185%3ATOC.

Expósito, R. G., de Bruijn, I., Postma, J., and Raaijmakers, J. M. (2017). Current Insights into the Role of Rhizosphere Bacteria in Disease Suppressive Soils. Frontiers in Microbiology 8. issn: 1664-302X. doi: 10.3389/fmicb.2017.02529.

Frank, J. A., Reich, C. I., Sharma, S., Weisbaum, J. S., Wilson, B. A., and Olsen, G. J. (2008). Critical Evaluation of Two Primers Commonly Used for Amplification of Bacterial 16S rRNA Genes. Applied and Environmental Microbiology 74.8, pp. 2461–2470. doi: 10.1128/AEM.02272-07.

Gloor, G. B., Macklaim, J. M., Pawlowsky-Glahn, V., and Egozcue, J. J. (2017). Microbiome Datasets Are Compositional: And This Is Not Optional. Frontiers in Microbiology 8. issn: 1664-302X. doi: 10.3389/fmicb.2017.02224.

Gu (2022). Complex Heatmap Visualization. iMeta 1.3, e43. issn: 2770-596X, 2770-596X. doi: 10.1002/imt2.43.

Gu, Banerjee, S., Dini-Andreote, F., Xu, Y., Shen, Q., Jousset, A., and Wei, Z. (2022). Small Changes in Rhizosphere Microbiome Composition Predict Disease Outcomes Earlier than Pathogen Density Variations. The ISME Journal 16.10, pp. 2448–2456. issn: 1751-7362. doi: 10.1038/s41396-022-01290-z.

Hajji, L., Hlaoua, W., Regaieg, H., and Horrigue-Raouani, N. (2017). Biocontrol Potential of Verticillium Leptobactrum and Purpureocillium Lilacinum Against Meloidogyne Javanica and Globodera Pallida on Potato (Solanum Tuberosum). American Journal of Potato Research 94.2, pp. 178–183. issn: 1874-9380. doi: 10.1007/s12230-016-9554-0.

Harkes, P., Suleiman, A. K. A., van den Elsen, S. J. J., de Haan, J. J., Holterman, M., Kuramae, E. E., and Helder, J. (2019). Conventional and Organic Soil Management as Divergent Drivers of Resident and Active Fractions of Major Soil Food Web Constituents. Scientific Reports 9.1, p. 13521. issn: 2045-2322. doi: 10.1038/s41598-019-49854-y.

Henry, A. W. (1931). The Natural Microflora of the Soil in Relation to the Foot-Rot Problem of Wheat. Canadian Journal of Research 4.1, pp. 69–77. issn: 1923-4287. doi: 10.1139/cjr31-006.

Hu, W., Samac, D. A., Liu, X., and Chen, S. (2017). Microbial Communities in the Cysts of Soybean Cyst Nematode Affected by Tillage and Biocide in a Suppressive Soil. Applied Soil Ecology 119, pp. 396–406. issn: 0929-1393. doi: 10.1016/j.apsoil.2017.07.018.

Indarti, S., Widianto, D., Mulyadi, M., and Widada, J. (2021). Potato Cyst Nematode-Infected Soil as a Source of Egg and Cyst Parasitic Fungi as Potential Biocontrol Agents. Archives of Phytopathology and Plant Protection 54.13–14, pp. 850–869. issn: 0323-5408. doi: 10.1080/03235408.2020.1857152.

Jones (2024). Characterisation of Microbial Communities Associated with Potato Cyst Nematodes in a Suppressive Soil and the Bicontrol Potential of Selected Isolates. doi: 10.17869/enu.2024.3789821.

Jones, et al. (2013). Top 10 Plant-Parasitic Nematodes in Molecular Plant Pathology. Molecular Plant Pathology 14.9, pp. 946–961. issn: 1364-3703. doi: 10.1111/mpp.12057.

Kauserud, H. (2023). ITS Alchemy: On the Use of ITS as a DNA Marker in Fungal Ecology. Fungal Ecology 65, p. 101274. issn: 1754-5048. doi: 10.1016/j.funeco.2023.101274.

Kerry, B., Davies, K., and Esteves, I. (2009). Work Undertaken between: 1st January 2006 and 31st December 2008. Final Report.

Lauber, C. L., Strickland, M. S., Bradford, M. A., and Fierer, N. (2008). The Influence of Soil Properties on the Structure of Bacterial and Fungal Communities across Land-Use Types. Soil Biology and Biochemistry. Special Section: Enzymes in the Environment 40.9, pp. 2407–2415. issn: 0038-0717. doi: 10.1016/j.soilbio.2008.05.021.

Li, J., Zou, C., Xu, J., Ji, X., Niu, X., Yang, J., Huang, X., and Zhang, K.-Q. (2015). Molecular Mechanisms of Nematode-Nematophagous Microbe Interactions: Basis for Biological Control of Plant-Parasitic Nematodes. Annual Review of Phytopathology 53.Volume 53, 2015, pp. 67–95. issn: 0066-4286, 1545-2107. doi: 10.1146/annurev-phyto-080614-120336.

Lin, H. and Peddada, S. D. (2020). Analysis of Compositions of Microbiomes with Bias Correction. Nature Communications 11.1, p. 3514. issn: 2041-1723. doi: 10.1038/s41467-020-17041-7.

Love, M. I., Huber, W., and Anders, S. (2014). Moderated estimation of fold change and dispersion for RNA-seq data with DESeq2. Genome Biology 15 (12), p. 550. doi: 10.1186/s13059-014-0550-8.

Manzanilla-Ĺopez, R. H., Esteves, I., and Devonshire, J. (2017). “Biology and Management of Pochonia Chlamydosporia and Plant-Parasitic Nematodes”. In: Perspectives in Sustainable Nematode Management Through Pochonia Chlamydosporia Applications for Root and Rhizosphere Health. Ed. by R. H. Manzanilla-Ĺopez and L. V. Lopez-Llorca. Cham: Springer International Publishing, pp. 47–76. isbn: 978-3-319-59224-4. doi: 10.1007/978-3-319-59224-4_3.

Martin, M. (2011). Cutadapt Removes Adapter Sequences from High-Throughput Sequencing Reads. EMBnet.journal 17.1, pp. 10–12. issn: 2226-6089. doi: 10.14806/ej.17.1.200.

Martino, C., Morton, J. T., Marotz, C. A., Thompson, L. R., Tripathi, A., Knight, R., and Zengler, K. (2019). A Novel Sparse Compositional Technique Reveals Microbial Perturbations. mSystems 4.1, 10.1128/msystems.00016–19. doi: 10.1128/msystems.00016-19.

McMurdie, P. J. and Holmes, S. (2013). phyloseq: An R package for reproducible interactive analysis and graphics of microbiome census data. PLoS ONE 8.4, e61217. url: http://dx.plos.org/10.1371/journal.pone.0061217.

Mhatre, P. H., Divya, K., Venkatasalam, E., Watpade, S., Bairwa, A., and Patil, J. (2022). Management of Potato Cyst Nematodes with Special Focus on Biological Control and Trap Cropping Strategies. Pest Management Science 78.9, pp. 3746–3759. issn: 1526-4998. doi: 10.1002/ps.7022.

Nagachandrabose, S. (2020). Management of Potato Cyst Nematodes Using Liquid Bioformulations of Pseudomonas Fluorescens, Purpureocillium Lilacinum and Trichoderma Viride. Potato Research 63.4, pp. 479–496. issn: 1871-4528. doi: 10.1007/s11540-020-09452-2.

Nordbring-Hertz, B. (2004). Morphogenesis in the Nematode-Trapping Fungus Arthrobotrys Oligospora - an Extensive Plasticity of Infection Structures. Mycologist 18.3, pp. 125–133. issn: 0269-915X, 1474-0605. doi: 10.1017/S0269915X04003052.

Nyang’au, M. N., Akutse, K. S., Fathiya, K., Charimbu, M. K., and Haukeland, S. (2023). Biodiversity and Efficacy of Fungal Isolates Associated with Kenyan Populations of Potato Cyst Nematode (*Globodera* Spp.) Biological Control 186, p. 105328. issn: 1049-9644. doi: 10.1016/j.biocontrol.2023.105328.

Oksanen, J. et al. (2022). vegan: Community Ecology Package. R package version 2.6-4. url: https://CRAN.R-project.org/package=vegan.

Ossowicki, A., Tracanna, V., Petrus, M. L. C., van Wezel, G., Raaijmakers, J. M., Medema, M. H., and Garbeva, P. (2020). Microbial and Volatile Profiling of Soils Suppressive to Fusarium Culmorum of Wheat. Proceedings of the Royal Society B: Biological Sciences 287.1921, p. 20192527. doi: 10.1098/rspb.2019.2527.

Penton, C. R., Gupta, V. V. S. R., Tiedje, J. M., Neate, S. M., Ophel-Keller, K., Gillings, M., Harvey, P., Pham, A., and Roget, D. K. (2014). Fungal Community Structure in Disease Suppressive Soils Assessed by 28S LSU Gene Sequencing. PLOS ONE 9.4, e93893. issn: 1932-6203. doi: 10.1371/journal.pone.0093893.

Powell, J. R., Karunaratne, S., Campbell, C. D., Yao, H., Robinson, L., and Singh, B. K. (2015). Deterministic Processes Vary during Community Assembly for Ecologically Dissimilar Taxa. Nature Communications 6.1, p. 8444. issn: 2041-1723. doi: 10.1038/ncomms9444.

Quast, C., Pruesse, E., Yilmaz, P., Gerken, J., Schweer, T., Yarza, P., Peplies, J., and Glöckner, F. O. (2012). The SILVA Ribosomal RNA Gene Database Project: Improved Data Processing and Web-Based Tools. Nucleic acids research 41.D1, pp. D590–D596.

Quensen, J. (2020). QsRutils: R Functions Useful for Community Ecology. R package version 0.1.5. url: https://github.com/jfq3/QsRutils.

RCoreTeam (2013). R: A Language and Environment for Statistical Computing. Foundation for Statistical Computing, Vienna, Austria.

Rivers, A. R., Weber, K. C., Gardner, T. G., Liu, S., and Armstrong, S. D. (2018). ITSxpress: Software to Rapidly Trim Internally Transcribed Spacer Sequences with Quality Scores for Marker Gene Analysis. F1000Research 7.

Rognes, T., Flouri, T., Nichols, B., Quince, C., and Mahé, F. (2016). VSEARCH: A Versatile Open Source Tool for Metagenomics. PeerJ 4, e2584.

Rousk, J., Bååth, E., Brookes, P. C., Lauber, C. L., Lozupone, C., Caporaso, J. G., Knight, R., and Fierer, N. (2010). Soil Bacterial and Fungal Communities across a pH Gradient in an Arable Soil. The ISME Journal 4.10, pp. 1340–1351. issn: 1751-7362, 1751-7370. doi: 10.1038/ismej.2010.58.

Sankaranarayanan, C. and Hari, K. (2021). Integration of Arbuscular Mycorrhizal and Nematode Antagonistic Fungi for the Biocontrol of Root Lesion Nematode Pratylenchus Zeae Graham, 1951 on Sugarcane. Sugar Tech 23.1, pp. 194–200. issn: 0974-0740. doi: 10.1007/s12355-020-00876-1.

Santos, M. C. V. dos, Esteves, I., Kerry, B., and Abrantes, I. (2013). Biology, Growth Parameters and Enzymatic Activity of Pochonia Chlamydosporia Isolated from Potato Cyst and Root-Knot Nematodes. doi: 10.1163/15685411-00002695.

Scholler, M., Hagedorn, G., and Rubner, A. (1999). A Reevaluation of Predatory Orbiliaceous Fungi. II. A New Generic Concept.

Scholler, M., Hagedorn, G., and Rubner, A. (2000). Arthrobotrys Oudemansii Nom. Nov., a New Name for a Nematode-Trapping Fungus to Avoid Homonymy.

Seinhorst, J. W. (1964). Methods for the Extraction of Heterodera Cysts from Not Previously Dried Soil Samples.

Singh, S., Pandey, R. K., and Goswami, B. (2013). Bio-Control Activity of Purpureocillium Lilacinum Strains in Managing Root-Knot Disease of Tomato Caused by Meloidogyne Incognita. Biocontrol Science and Technology 23.12, pp. 1469–1489. issn: 0958-3157. doi: 10.1080/09583157.2013.840770.

Soliman, M., El-Deriny, M., Ibrahim, D., Zakaria, H., and Ahmed, Y. (2021). Suppression of Root-knot Nematode Meloidogyne Incognita on Tomato Plants Using the Nematode Trapping Fungus Arthrobotrys Oligospora Fresenius. Journal of Applied Microbiology 131.5, pp. 2402–2415. issn: 1364-5072. doi: 10.1111/jam.15101.

Stirling, G. R. (2014). *Biological Control of Plant-parasitic Nematodes*, *2nd Edition: Soil Ecosystem Management in Sustainable Agriculture*. CABI. isbn: 978-1-78064-415-8.

Sun, R., Dsouza, M., Gilbert, J. A., Guo, X., Wang, D., Guo, Z., Ni, Y., and Chu, H. (2016). Fungal Community Composition in Soils Subjected to Long-Term Chemical Fertilization Is Most Influenced by the Type of Organic Matter. Environmental Microbiology 18.12, pp. 5137–5150. issn: 1462-2920. doi: 10.1111/1462-2920.13512.

Tedersoo, L. and Anslan, S. (2019). Towards PacBio-based Pan-Eukaryote Metabarcoding Using Full-Length ITS Sequences. Environmental Microbiology Reports 11.5, pp. 659– 668. issn: 1758-2229. doi: 10.1111/1758-2229.12776.

Teklu, M. G., Beniers, J. E., Been, T. H., Schomaker, C. H., and Molendijk, L. P. G. (2018). Methods for the Estimation of Partial Resistance in the Glasshouse of Potato Cultivars/Genotypes against Potato Cyst Nematodes & Root-Knot Nematodes.

Thakur, M. P. and Geisen, S. (2019). Trophic Regulations of the Soil Microbiome. Trends in Microbiology 27.9, pp. 771–780. issn: 0966-842X. doi: 10.1016/j.tim.2019.04.008.

Topalovíc, O., Hussain, M., and Heuer, H. (2020). Plants and Associated Soil Microbiota Cooperatively Suppress Plant-Parasitic Nematodes. Frontiers in Microbiology 11. issn: 1664-302X. doi: 10.3389/fmicb.2020.00313.

Velvis, H. and Kamp, P. (1996). Suppression of Potato Cyst Nematode Root Penetration by the Endoparasitic Nematophagous fungusHirsutella Rhossiliensis. European Journal of Plant Pathology 102.2, pp. 115–122. issn: 1573-8469. doi: 10.1007/BF01877097.

Venables, W. N. and Ripley, B. D. (2002). Modern Applied Statistics with S. Fourth. ISBN 0-387-95457-0. New York: Springer. url: https://www.stats.ox.ac.uk/pub/MASS4/.

Watson, T. T., Strauss, S. L., and Desaeger, J. A. (2020). Identification and Characterization of Javanese Root-Knot Nematode (*Meloidogyne Javanica*) Suppressive Soils in Florida. Applied Soil Ecology 154, p. 103597. issn: 0929-1393. doi: 10.1016/j.apsoil.2020.103597.

Wei, Z., Gu, Y., Friman, V.-P., Kowalchuk, G. A., Xu, Y., Shen, Q., and Jousset, A. (2019). Initial Soil Microbiome Composition and Functioning Predetermine Future Plant Health. Science Advances 5.9, eaaw0759. issn: 2375-2548. doi: 10.1126/sciadv.aaw0759.

Weller, D., Letourneau, M., and Yang, M. (2022). “Classification, Discovery, and Microbial Basis of Disease-Suppressive Soils”. In: pp. 457–465. isbn: 978-1-119-76262-1. doi: 10. 1002/9781119762621.ch36.

Wernet, V. and Fischer, R. (2023). Establishment of Arthrobotrys Flagrans as Biocontrol Agent against the Root Pathogenic Nematode Xiphinema Index. Environmental Microbiology 25.2, pp. 283–293. issn: 1462-2920. doi: 10.1111/1462-2920.16282.

Wickham, H. (2016). ggplot2: Elegant Graphics for Data Analysis. Springer-Verlag New York. isbn: 978-3-319-24277-4. url: https://ggplot2.tidyverse.org.

Yu, Q. (2011). Soybean Cyst Nematode (Heterodera Glycines Ichinohe). HA El-Shemy (éd.). Soybean Physiology and Biochemistry. InTech, New York, NY, É.-U, pp. 461–474.

Zhou, D., Feng, H., Schuelke, T., De Santiago, A., Zhang, Q., Zhang, J., Luo, C., and Wei, L. (2019). Rhizosphere Microbiomes from Root Knot Nematode Non-infested Plants Suppress Nematode Infection. Microbial Ecology 78.2, pp. 470–481. issn: 1432-184X. doi: 10.1007/s00248-019-01319-5.

