## Supplementary material for "Patchy distribution of potato cyst nematodes within single arable fields reveals local disease suppressiveness mediated by disparate microbial communities"

### 1 Supplementary material

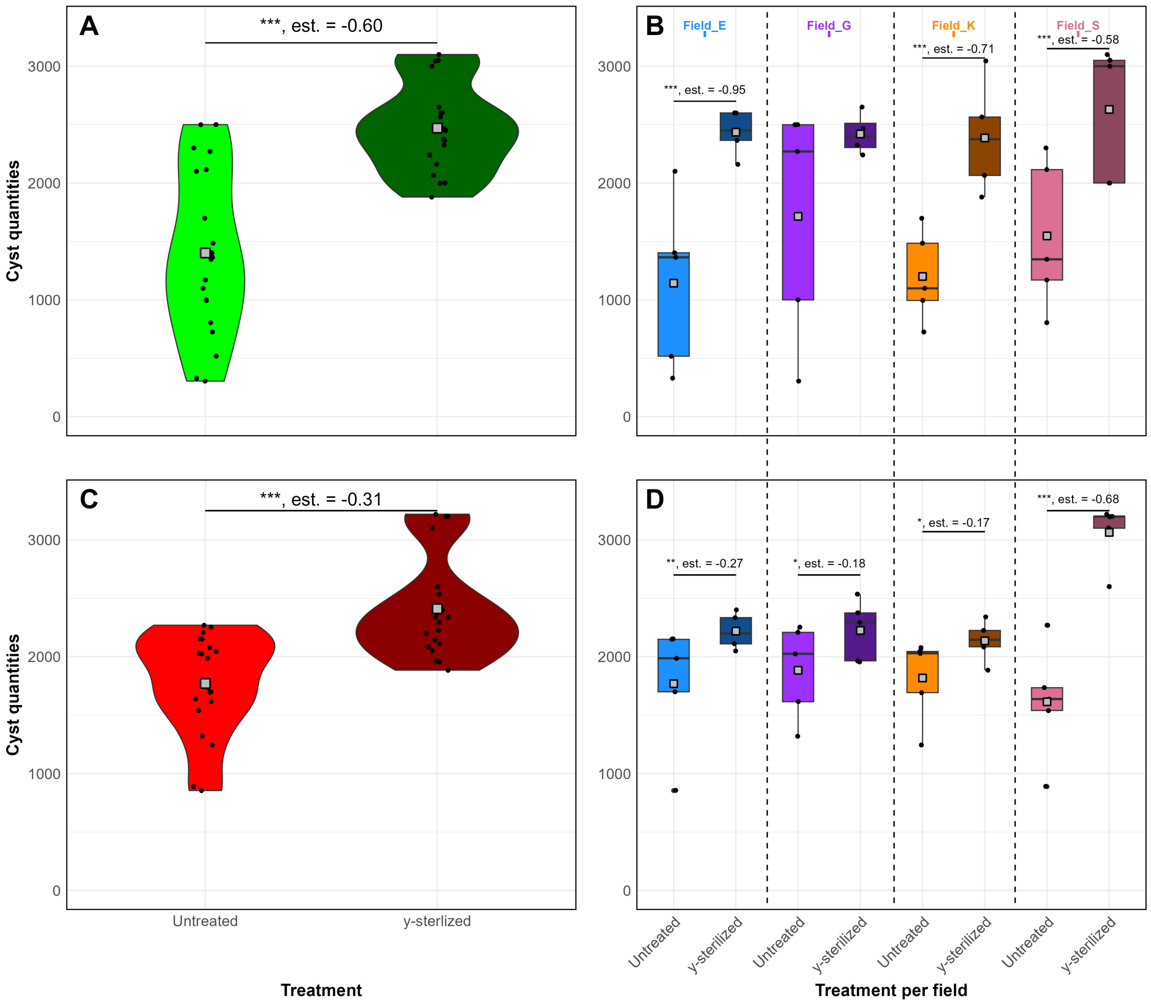


Supplementary Figure S1: Number of potato cyst nematode cysts per pot quantified at 16 weeks post-inoculation on fully susceptible potato plants. A) Cyst counts of the untreated and sterilized soil are shown for the putatively suppressive parts aggregated across all fields (n=20) and B) per field (n=5), as well as for the putatively conducive patches aggregated across C) all fields and D) per field. * (P <0.05) and ** (P <0.01), *** (P<0.001) indicate statistical significant difference as determined by a GLM with a Negative Binomial distribution.

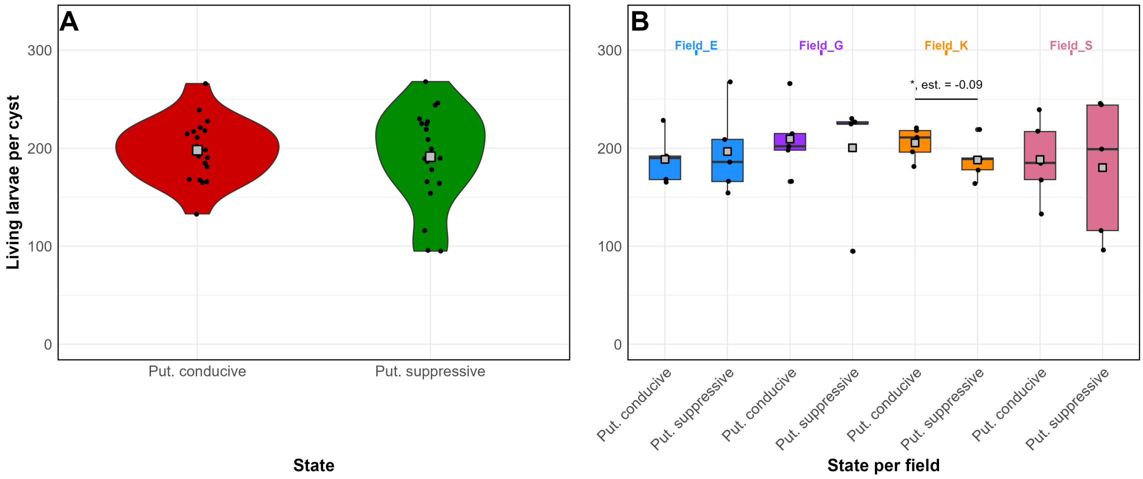


Supplementary Figure S2: Number of potato cyst nematode living larvae per cyst per pot quantified at 16 weeks post-inoculation on fully susceptible potato plants in putatively conducive and suppressive soil. A) Living larvae counts are shown for untreated soil aggregated across all fields ( n=20) and B) per field (n=5). * ($P < 0.05$) indicates statistical significant difference as determined by a GLM with a Negative Binomial distribution. Put. = putatively.

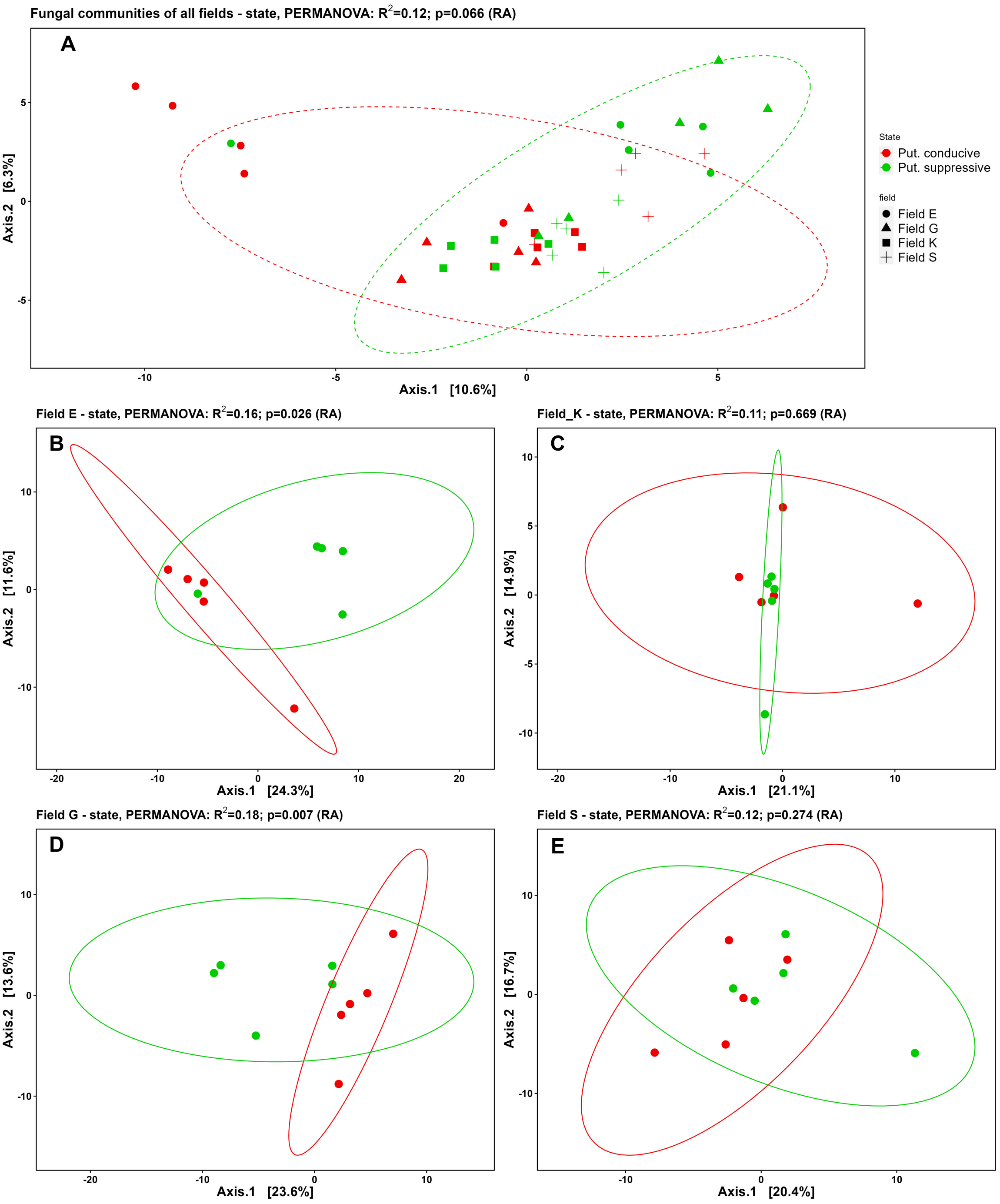


Supplementary Figure S3: PCoA plots based on the robust Aitchison distance of the fungal communities in rhizosphere samples collected from potato plants 13 weeks after inoculation with potato cyst nematodes. A) Fungal communities in rhizosphere of plants exposed to putatively conducive (red) or suppressive (green) soil from all four field locations and B-E) at individual field location level (n=5). B = field E, C = field K, D = field G, and E = field S. Next to A), symbols for the individual field locations are presented. Figure headers include the PERMANOVA result of factor 'state' (= putatively supressive or conducive). Put. = putatively.

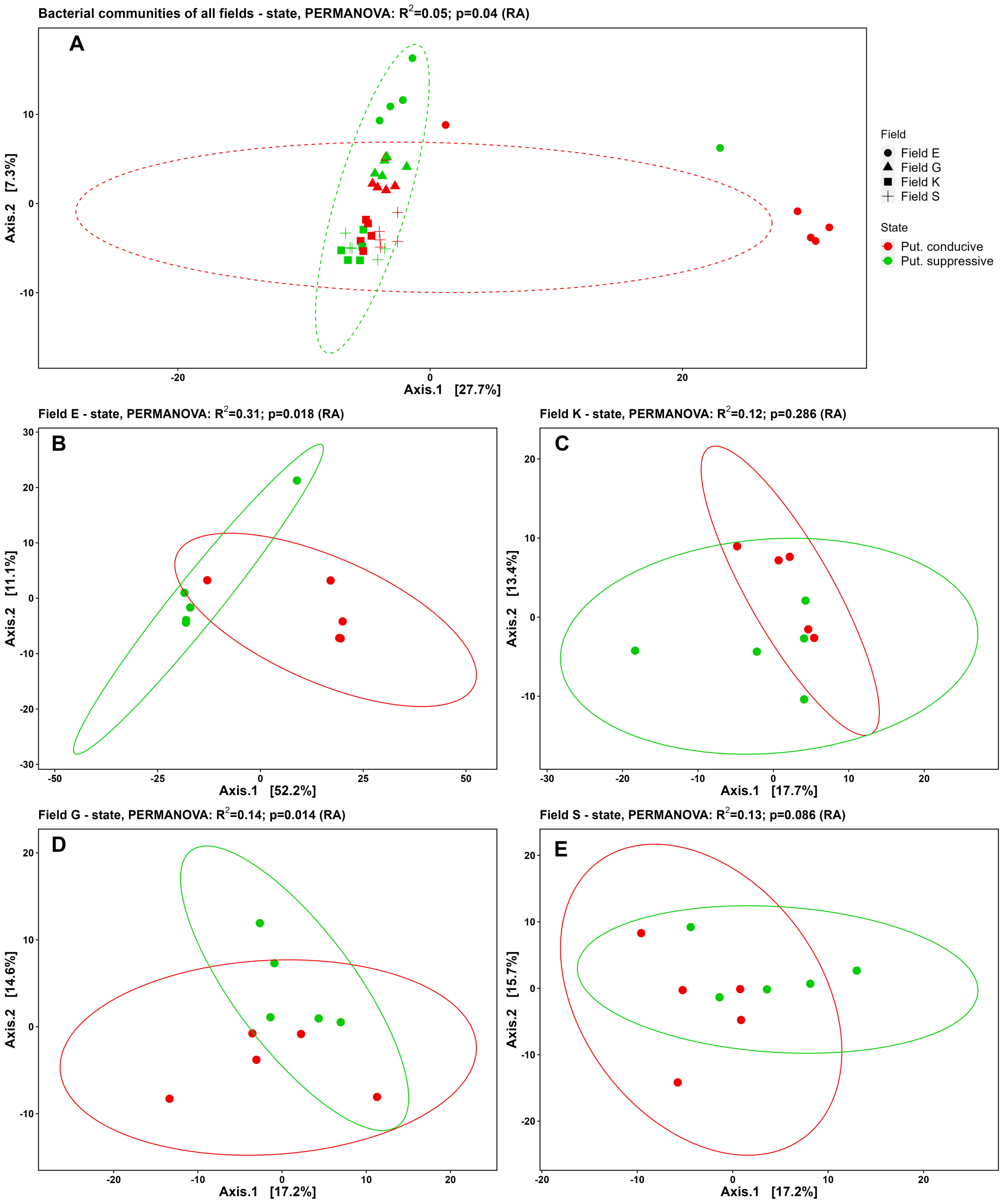


Supplementary Figure S4: PCoA plots based on the robust Aitchison distance of the bacterial communities in rhizosphere samples collected from potato plants 13 weeks after inoculation with potato cyst nematodes. A) Bacterial communities in rhizosphere of plants exposed to putatively conducive (red) or suppressive (green) soil from all four field locations, and B-E) at individual field location level (n=5). B = field E, C = field K, D = field G, and E = field S. Next to A), symbols for the individual field locations are presented. Figure headers include the PERMANOVA result of factor state (= putatively supressive or conducive). Put. = putatively.

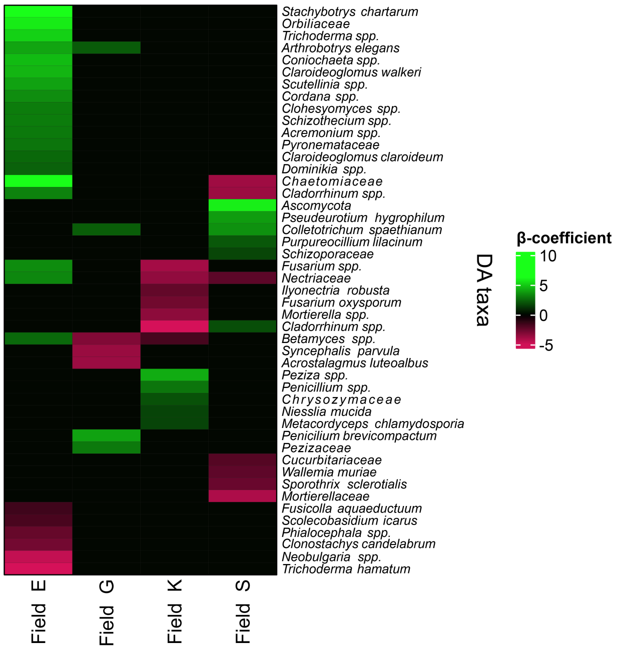


Supplementary Figure S5: Fungal differential abundance analysis (ANCOM-BC) per field, based on the Holm adjusted P-value with a significant threshold of 0.05. Taxa are shown in the rows and fields in the columns. Only species, genera, or families with a β-coefficient lower than -1.5 and larger than 1.5 are shown. Taxa with a positive β-coefficient (= green) are more abundant in the suppressive part as compared to the conducive part, while taxa with a negative (= red) β-coefficient are less abundant in the suppressive part as compared to the conducive part.

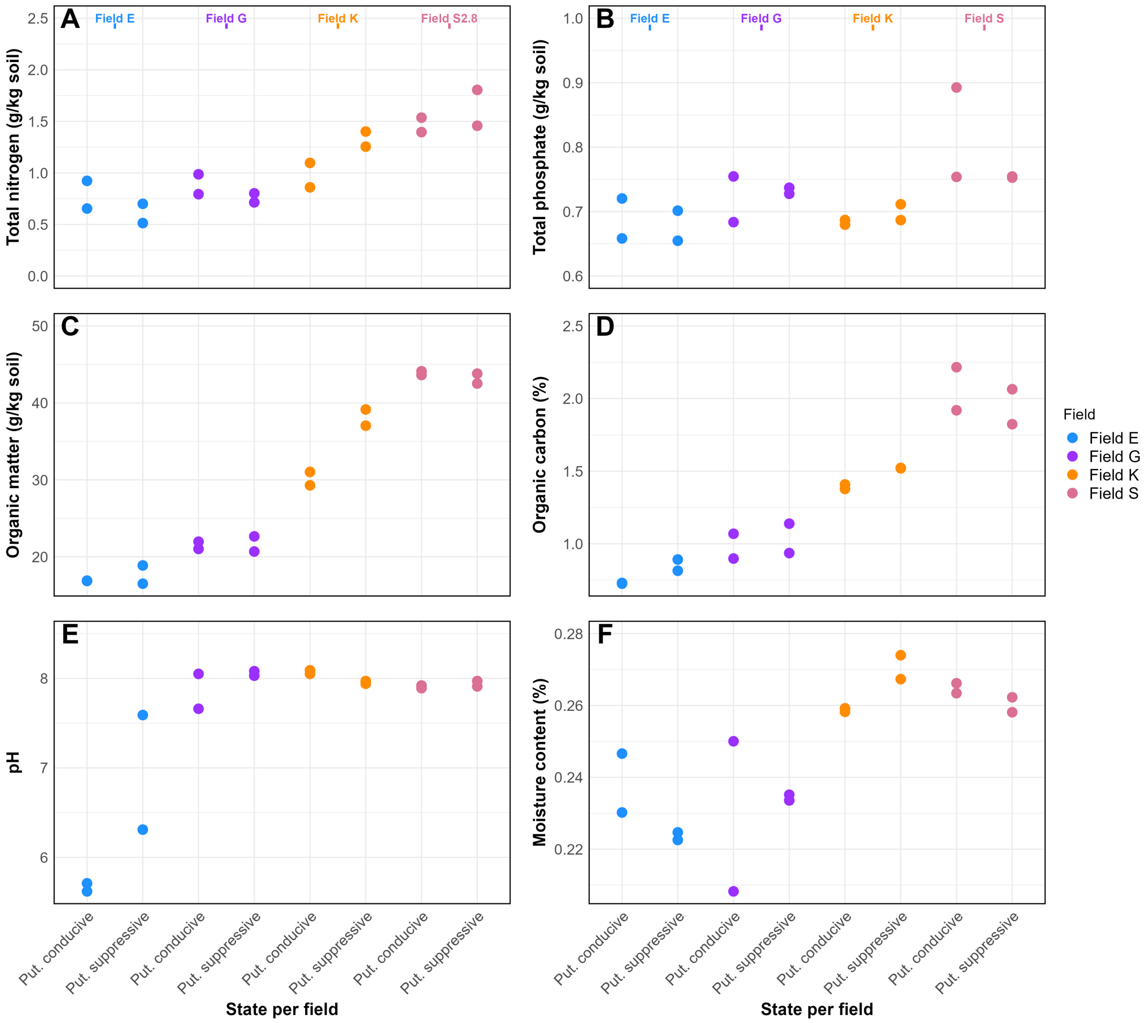


Supplementary Figure S6: Chemical analyses of two plots per field per state (=putatively conducive or putatively suppressive). A) Total nitrogen, B) Total phosphate, C) Organic matter, D) Organic carbon. E) pH, and F) moisture content. Colors indicate fields (blue = field E, purple = field G, orange = field K, pink = field S). Put.=putatively.
